# Bioengineered algal lipids enriched in structured medium- and long-chain triacylglycerols, linoleate, and *sn*-2 palmitate for human milkfat substitutes

**DOI:** 10.64898/2026.05.15.725531

**Authors:** Jon Y-T. Lin, Marco A. Dueñas, Suzanne M. Kosina, Anthony T. Iavarone, Kenzo Khoo, Carrie D. Nicora, Samuel O. Purvine, Trent R. Northen, Jeffrey L. Moseley, Sabeeha S. Merchant

**Affiliations:** California Institute for Quantitative Biosciences, University of California, Berkeley, Berkeley, CA 94720, USA; Department of Plant and Microbial Biology, University of California, Berkeley, Berkeley, CA 94720, USA; Department of Molecular and Cell Biology, University of California, Berkeley, Berkeley, CA 94720, USA; Biological Sciences Division, Pacific Northwest National Laboratory, United States Department of Energy, Richland, WA 99352, USA; Environmental Molecular Sciences Laboratory, Pacific Northwest National Laboratory, United States Department of Energy, Richland, WA 99352, USA; Environmental Genomics and Systems Biology, Lawrence Berkeley National Laboratory Berkeley, Berkeley, CA 94720, USA; Department of Chemistry, University of California, Berkeley, Berkeley, CA 94720, USA; Joint Genome Institute, Lawrence Berkeley National Laboratory, Berkeley, CA 94720, USA

## Abstract

Triacylglycerols (TAGs) in human milkfat (HMF) provide over 50% of the calories for infant nutrition. Their unique architecture, featuring palmitic acid predominantly esterified at the *sn*-2 position, facilitates absorption as 2-palmitoyl monoacylglycerol after hydrolysis of the *sn*-1 and *sn*-3 fatty acids by gut lipases. Conventional vegetable oil-based infant formulas mimic the HMF fatty acid profile but lack its distinct regioisomeric structure, abundant medium- and long-chain triglycerides (MLCTs), and oleic-to-linoleic acid ratios that vary with maternal diet and geography. We have engineered the oleaginous green alga *Auxenochlorella* for biosynthesis of TAGs enriched in MLCTs, *sn*-2 palmitate, and linoleate via heterologous expression of acyl-acyl-carrier-protein (ACP) thioesterases, palmitate-specific lysophosphatidic acid acyltransferases, and lysophosphatidylcholine acyltransferase. These structured algal lipids replicate both the major fatty acid proportions and regioisomeric composition of authentic HMF, providing single-source bioengineered alternatives to plant-derived HMF substitutes for infant formula.

## Introduction

Human milk is a complex biofluid that provides infants with complete nutrition for up to six months postpartum^1^. The fat component, composed primarily of triacylglycerols (TAGs), provides half of the energy for growth, and also includes bioactive and structural phospholipids, sphingolipids, sterols and fatty acids with signaling and developmental roles^2^. Palmitate (C16:0), oleate (C18:1 n-9) and linoleate (C18:2 n-6) are the most abundant acyl groups in human milkfat (HMF), and their regioisomeric positioning in TAGs is important for infant lipid assimilation^3^. TAG hydrolysis by gastric and intestinal lipases produces free fatty acids from the glycerol *sn*-1 and *sn*-3 positions, along with 2-monoacylglycerol (2-MAG)^4^. Saturated, medium-chain fatty acids (MCFA, C6:0–12:0) or unsaturated, long-chain fatty acids hydrolyzed from the TAG *sn*-1 and *sn*-3 positions by gastric and intestinal lipases are readily absorbed by enterocytes^4,5^, but palmitate, highly enriched at *sn*-2, is absorbed as 2-palmitoylglycerol, mitigating its poor solubility as the free acid6. In contrast, vegetable oil blends commonly used as lipid sources in infant formulas are typically enriched in unsaturated long-chain fatty acids at *sn*-2, while palmitate is preferentially found at *sn*-1 or *sn*-3^7^. The lipases thus release palmitic acid which forms non-absorbable calcium soaps, leading to infant intestinal distress, reduced caloric intake, and calcium deficiency in extreme cases^8,9^.

Additional defining characteristics of HMF are the substantial proportions of MCFA (6−16% of total FA)^10,11^ and linoleate (C18:2 n-6) (15−25% of total FA)^10,12^. These levels are shaped by maternal diet and vary between geographic regions. Linoleic acid and α-linolenic acid (C18:3 n-3) are essential fatty acids for mammals^13–15^, and U.S. regulations require linoleate supplementation in formulas^16,17^. MCFA are a key energy source due to their direct transport from enterocytes to the liver via the hepatic portal vein^5,18^, as well as bypass of the carnitine shuttle for import into mitochondria for oxidation^19,20^. MCFA in plant-based formulas are mainly derived from coconut or palm kernel oils rich in trisaturated medium-chain TAGs (MCTs), in contrast with the mixed medium- and long-chain triglycerides (MLCTs) found in HMF^21^. Clinical data have shown that MLCTs provide improved health benefits to infants through as yet unclear mechanisms^21–23^.

HMF substitutes with high *sn*-2 palmitate for infant formula are produced commercially by enzymatic interesterification of fractionated lipids and free fatty acids derived from chemical hydrolysis or lipase treatment of natural oils^24,25^. Metabolic engineering for 1-oleoyl-2-palmitoyl-3-oleoylglycerol (OPO) enriched triglycerides has been reported in Arabidopsis^26,27^, in the oleaginous yeast *Yarrowia lipolytica*^28^ and in *Prototheca wickerhamii*, a non-photosynthetic trebouxiophyte alga^29,30^. *sn*-2 palmitate selectivity was achieved by expressing plant, algal or human lysophosphatidic acid acyltransferases (LPAATs) with specificity for palmitoyl-coenzyme A substrates^26–30^. However, neither enzymatic nor synthetic biology approaches have produced single-source oils that mimic both the complex fatty acid composition of HMF and the regioisomeric *sn*-2 positioning of palmitic acid.

*Auxenochlorella* spp. are emerging reference organisms for discovery biology and biotechnology^31,32^. Homologous recombination in the nuclear genome facilitates effective gene targeting and reproducible transgene expression^31–33^. The synthetic biology toolkit for these species includes well-annotated genome assemblies, neutral landing sites for integration, biosafe transformation markers, and protein targeting to intracellular compartments^31,32,34^. TAG accumulation is maximized under heterotrophic cultivation with reduced carbon sources^35^, but unlike closely-related *Prototheca*^36,37^, *Auxenochlorella* retain the capability for reversible switching between photoautotrophic and heterotrophic growth^38^. These features offer advantages for lipid engineering over obligate heterotrophic microbial platforms, due to an inherent capacity for robust cofactor biosynthesis^39^ and intrinsic accumulation of photoprotective carotenoids and tocopherols that shield lipids from oxidative damage^40^. In this study, we build on prior *sn*-2 palmitate bioengineering efforts and describe systematic, multi-step engineering of the hybrid strain, *Auxenochlorella protothecoides x symbiontica* UTEX 250-A (hereafter referred to as *Auxenochlorella*) to produce TAGs suitable as HMF substitutes. This approach yielded single source lipids that replicate the MCFA, linoleic acid, *sn*-2 palmitate and MLCT composition characteristic of HMF.

## Results

### Engineering *Auxenochlorella* lipids to match the major fatty acid composition of HMF

*Auxenochlorella* storage lipid fatty acid profiles are simpler than those of HMF, with oleic acid (C18:1 n-9) as the dominant species, linoleic and α-linolenic acids as the most abundant polyunsaturates, and palmitic, stearic (C18:0) and myristic (C14:0) acids as the major saturated fatty acids (Fig. 1A and S1). Compared to *Auxenochlorella* lipids, HMF has less oleate but more palmitate and MCFA (Fig. 1A). Since fatty acid chain length is dictated by the substrate specificity of thioesterases that cleave the elongating acyl group from acyl carrier protein (ACP), we introduced heterologous plant thioesterases to modify *Auxenochlorella* fatty acid composition (Fig. 1B and Table S1). A codon-optimized gene encoding CwFATB2, a well-characterized fatty acyl-ACP thioesterase from *Cuphea wrightii* with selectivity for C10:0–C14:0^41^ was targeted to two distinct loci. *CwFATB2* was integrated at either allele 2 of the *Fatty Acid Biosynthesis 2A* (*FAB2A-2*) locus, encoding stearoyl-ACP desaturase, causing elevation of C18:0 at the expense of C18:1 n-9, or at the homozygous *DAO1* locus, encoding D-amino acid oxidase 1, which is not required for fatty acid biosynthesis or for growth under oil-producing conditions (Fig. 1A, C–D). Lauric (C12:0) and myristic acids increased in the modified strains, ranging between 3-21% of total fatty acids. The highest accumulation occurred in strains JL15 and JL21, in which the heterologous plant plastid-targeting transit peptide is replaced with the *Auxenochlorella* FAB2A plastid transit peptide (Fig. 1A, C–D)^42^. Untargeted proteomic analysis indicated that the endogenous plastid targeting sequence improves the accumulation of the introduced thioesterase by 3.5-fold (Table S2), consistent with more efficient import and processing of the foreign protein. Accordingly, stearic acid levels increased to 14.6–16.2% in heterozygous *FAB2A-1*/*fab2A-2* mutant strains (Fig. 1A, E), which exceeded the target range in HMF. Consequently, we pursued alternative strategies for achieving the HMF C18:0 target (4.8–6.2%, Fig. 1A).

**Figure 1.**
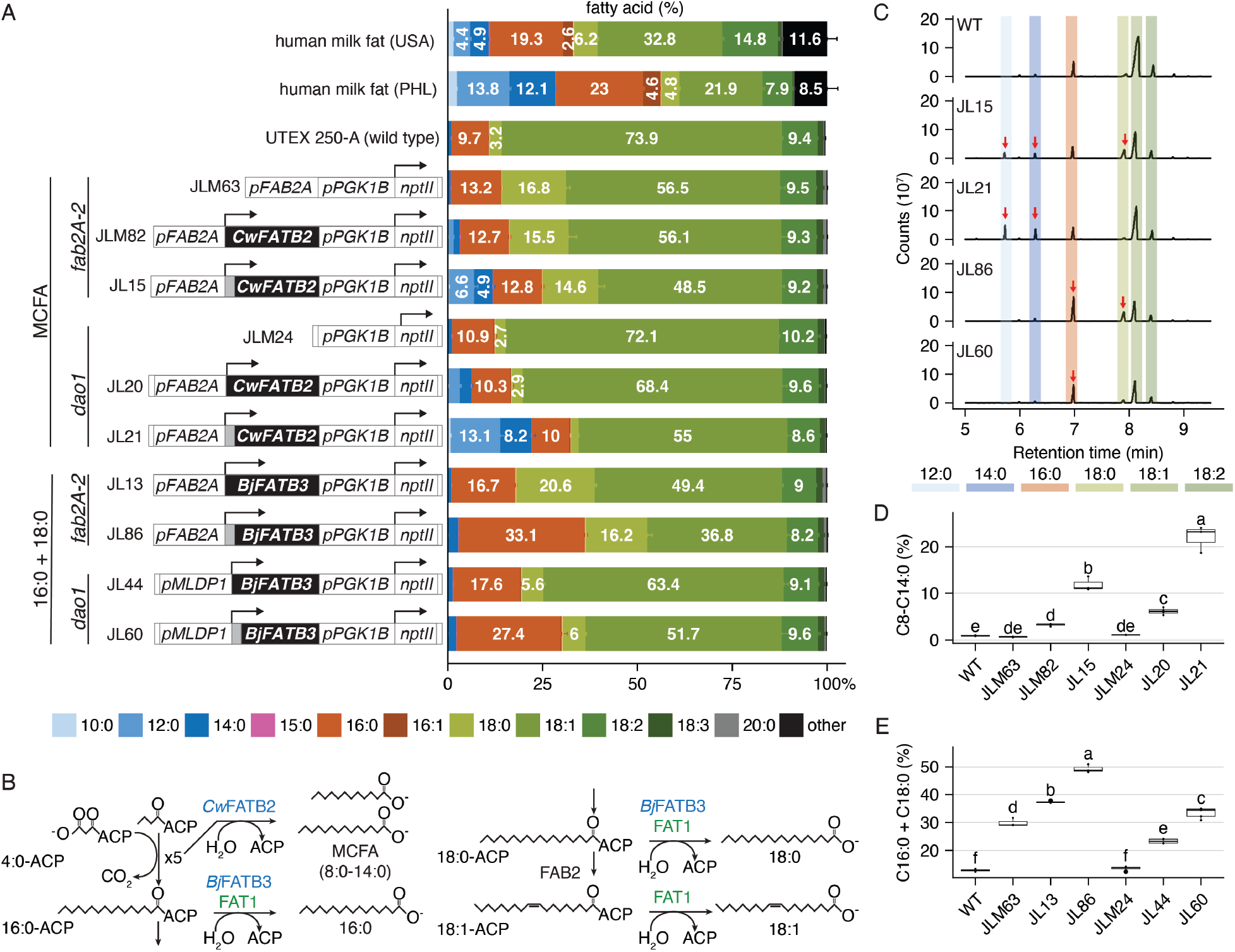
Heterologous thioesterases modify *Auxenochlorella* fatty acid chain length. **(A)** Fatty acid compositions (wt% of total fatty acids) of HMF from the United States (USA), Philippines (PHL)^3,10^, and heterotrophically grown *Auxenochlorella* wild type and engineered strains in this study. Schematics of the expression constructs are shown on the left. Constructs contain 5’ and 3’ DNA sequences targeting integration at the *FAB2A-2* (JLM63, JLM82, JL15, JL13, JL86) or *DAO1* (JLM24, JL20, JL21, JL44, JL60) loci by homologous recombination. Arrows represent the transcription start sites within promoters (*pFAB2A*, *pMLDP1*, *pPGK1B*) controlling downstream genes. *FAB2A* encodes stearoyl-ACP desaturase 2A, *MLDP1* encodes major lipid droplet protein 1, and *PGK1B* encodes phosphoglycerate kinase 1B. The *nptII* selection marker provides resistance to geneticin (G418). A sequence encoding the FAB2A-1 plastid transit peptide (grey) replaces the predicted CwFATB2 and BjFATB3 transit peptide-encoding regions in JL15, JL21, JL86, and JL60. **(B)** Fatty acid chain length engineering in *Auxenochlorella*. The endogenous FAT1 thioesterase is in green text. Heterologous thioesterases are in blue text. **(C)** Gas Chromatography-Mass Spectrometry (GC-MS) analysis of fatty acid methyl esters (FAMEs) prepared from wild-type and transgenic strain total lipids. The plots represent MS ion intensity versus GC retention time (min). Arrows indicate fatty acids that are significantly increased in abundance in the engineered strains compared to wild type. **(D)** C8–14:0 levels in strains expressing CwFATB2. **(E)** C16:0 plus C18:0 levels in strains expressing BjFATB3. n = 3-4 independent transformants of each transgenic strain. Refer to Table S1 for the strain details. n = 10 for the UTEX 250-A wild-type control. Error bars = SD. Letters represent Tukey Honest Significant Differences between strains (padj ≥0.05 for measurements with the same letter).

BjFATB3 from *Brassica juncea* was reported to increase C18:0 content in *E. coli*^43^. However, when expressed in *Auxenochlorella* with either its native or an algal plastid transit peptide, BjFATB3 increased both palmitic and stearic acid accumulation (Fig. 1A, C, E, compare strains JL44 and JL60 to JLM24 and wild type), suggesting in vivo substrate preference for both C16:0- and C18:0-ACP (Fig. 1B). The highest C16:0 levels were achieved in strains JL86 (33.1%) and JL60 (27.4%), compared to around 10% in the wild type (Fig. 1A, C, E). Stearic acid abundance doubled in strains expressing *BjFATB3* from the neutral *dao1* locus compared to the wild-type strain and the JLM24 transformation marker-only control, (Fig. 1A, C, E), bringing C18:0 into the HMF target range. We concluded that the HMF target amounts of C12:0, C14:0, C16:0 and C18:0 could be achieved via simultaneous expression of CwFATB2 and BjFATB3. We also noted that the JL13 and JL86 strains, expressing the *B. juncea* thioesterase gene in the *fab2A-2* mutant background, represent an intermediate engineering stage in the production of high-stearate structured fats such as those found in cocoa, mango and shea butters^44,45^.

### Enhanced C16:0 *sn-*2 regioisomeric positioning with heterologous LPAATs

C16:0 enrichment at the TAG *sn*-2 position is a key feature of HMF (Fig. 2A)^3^. Conversely, in common vegetable oils and in *Auxenochlorella*, the *sn*-2 position is predominantly esterified with unsaturated fatty acids (Fig. 2A). To alter C16:0 regioisomeric positioning in *Auxenochlorella*, we introduced ER-localized heterologous LPAATs with the desired palmitate substrate specificity (Fig. 2B–D)^46,47^. 23–25% of the TAGs in strains harboring *CrLPAAT2* from *Chlamydomonas* (JL27, JL73) or human *HsAGPAT1* (JL29) contained C16:0 at the *sn*-2 position, compared to the wild type with just 0.8–1.4% (Fig. 2D and Table 1). Total fatty acid profiles in the resulting strains did not change, with comparable C16:0 levels (10-11%) in wild type and in the LPAAT-expressors (Fig. 2D and Table 1). Accordingly, the fraction of total C16:0 at *sn*-2 increased from 2–5% in wild type to 72–75% in JL27 and JL29, within the HMF target range of 70–88% (Table 1)^3,48^. *sn*-2 incorporation of C14:0 and C18:0 increased modestly in the LPAAT-expressing strains, with a stronger effect in JL29 expressing HsAGPAT1, indicating that CrLPAAT2 is more selective for the C16:0 substrate than is the human enzyme (Table S3).

**Figure 2.**
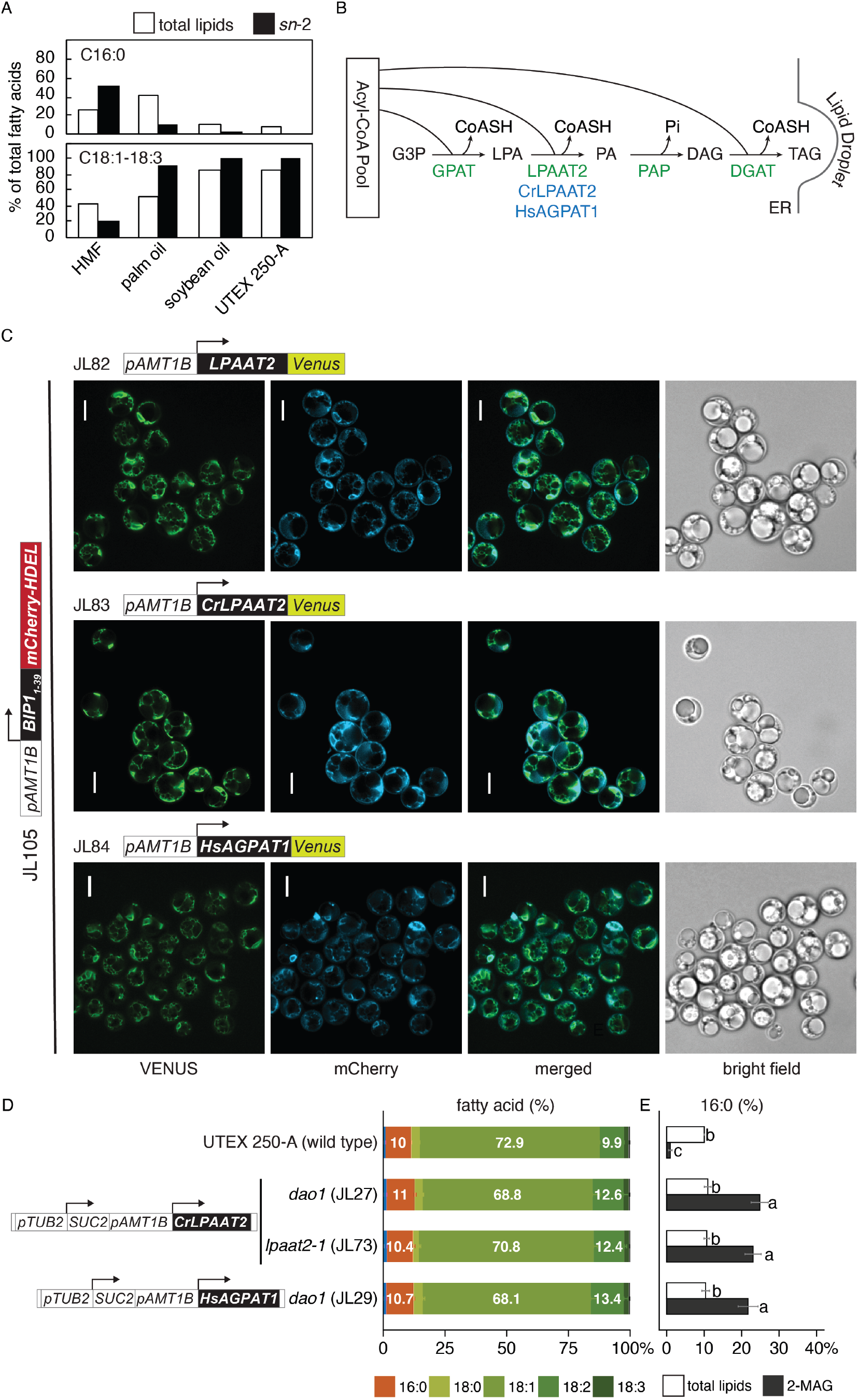
Heterologous ER lysophosphatidic acid acyltransferases enhance positioning of C16:0 at *sn*-2 in TAG. **(A)** C16:0 and unsaturated C18 (C18:1 n-9 + C18:2 n-6 + C18:3 n-3c) proportions of total fatty acids (white bars) and at *sn*-2 (black bars) in HMF ^3^, vegetable oils^3,100,101^, and in *Auxenochlorella* lipids. **(B)** Kennedy pathway engineering in *Auxenochlorella* for improved *sn*-2 palmitate biosynthesis. Endogenous *Auxenochlorella* enzymes are in green text; introduced genes are in blue text. G3P = glycerol-3-phosphate, LPA = lysophosphatidic acid, PA = phosphatidic acid, DAG = diacylglycerol. GPAT = glycerol-3-phosphate acyltransferase, LPAAT = lysophosphatidic acid acyltransferase, PAP = phosphatidic acid phosphatase, DGAT = diacylglycerol acyltransferase. **(C)** ER co-localization of C-terminal-Venus-fused LPAATs and N-terminal BIP1 signal-peptide fused to mCherry in lipid-producing cells (see Materials and Methods). Scale bar = 5 μm. **(D)** Fatty acid profiles (wt%) of strains expressing heterologous LPAATs. Only the most abundant fatty acids are shown (>1.5% of total). Error bars = SD. *SUC2* marker gene expression is controlled by the *C. reinhardtii TUB2* promoter (*pTUB2*), and the *Auxenochlorella AMMONIUM TRANSPORTER 1B* allele 1 promoter (*pAMT1B-1*) drives *CrLPAAT2* and *HsAGPAT1* gene transcription. **(E)** C16:0 percentage in total fatty acids (white bars) versus in 2-monoacylglycerol (2-MAG) (black bars) from *sn*-2 lipase assays. Letters indicate statistical groups determined by one-way ANOVA followed by Tukey’s post-hoc test. n = 4 independent transformants for JL27, JL29, and JL73, n = 3 independent cultures for the UTEX 250-A wild-type control. Errors bars = standard deviation.

**Table 1.** Regiospecific Distribution of Palmitate, MCFA, and Linoleate.

|  | HMF | *P.*<br>*moriformis*<sup>c</sup> | *A.*<br>*thaliana*<sup>d</sup> | *Y.*<br>*lipolytica*<sup>e</sup> | *Auxenochlorella* UTEX 250-A<sup>f</sup> |  |  |  |  |  |  |
| --- | --- | --- | --- | --- | --- | --- | --- | --- | --- | --- | --- |
|  |  |  |  |  | wild<br>type | JL27 | JL29 | JL52<br>+JL27 | JL88<br>+JL27 | JL88<br>+JL73 | JL116<br>+JL88<br>+JL27 |
| n | n/a | n/a | n/a | n/a | 6 | 4 | 4 | 3 | 3 | 4 | 4 |
| total<br>C16:0%<sup>ab</sup> | 25.0 | 31.5 | ~23.0 | ~24.0 | 10.00<br>±0.04 | 11.0<br>±0.9 | 10.7<br>±0.7 | 13.7<br>±0.7 | 20.1<br>±0.6 | 20.60<br>±0.14 | 20.7<br>±0.8 |
| *sn*-2<br>C16:0%<sup>ab</sup> | 51.0 | 69.3 | ~58.0 | ~50.0<sup>e</sup> | 1.0<br>±0.5 | 25<br>±2 | 23<br>±2 | 36<br>±2 | 43<br>±2 | 42<br>±3 | 45.2<br>±0.9 |
| %C16:0<br>at *sn*-2<sup>ab</sup> | 70.0<br>–<br>88.0 | 73.3 | ~84.0 | 69.1 | 3.2<br>±1.6 | 75.5<br>±0.9 | 72<br>±2 | 86.4<br>±1.2 | 71<br>±2 | 68<br>±5 | 73<br>±4 |
| total<br>MCFA%<sup>†</sup><br>ghi | 7.5–<br>16.5 | n/a | n/a | n/a | n.d. | n.d. | n.d. | 18.2<br>±1.0 | 12.4<br>±1.3 | 13.6<br>±0.1 | 14.5<br>±2.0 |
| *sn*-2<br>MCFA%<br>gh | 14.8<br>–<br>19.5 | n/a | n/a | n/a | n.d. | n.d. | n.d. | 5.5<br>±1.0 | 9.0<br>±0.4 | 11.0<br>±0.4 | 11.1<br>±1.9 |
| %MCFA<br>at *sn*-2<sup>gh</sup> | 35.5<br>–<br>49.3 | n/a | n/a | n/a | n.d. | n.d. | n.d. | 10.1<br>±2.5 | 24.2<br>±2.0 | 26.9<br>±1.1 | 26<br>±4 |
| total<br>C18:2%<br>aj | 13.0<br>–<br>21.4 | n/a | n/a | n/a | 9.90<br>±0.04 | 13<br>±2 | 13.4<br>±1.7 | 10.0<br>±0.7 | 11.1<br>±0.6 | 9.1<br>±0.3 | 25.4<br>±1.1 |
| *sn*-2<br>C18:2%<br>aj | 8.40<br>–<br>10.8 | n/a | n/a | n/a | 11.50<br>±0.16 | 9<br>±2 | 14<br>±2 | 7.8<br>±0.3 | 8.3<br>±0.5 | 6.4<br>±0.8 | 12.3<br>±0.4 |
| %C18:2<br>at *sn*-2%<br>aj | 16.8<br>–<br>21.5 | n/a | n/a | n/a | 38.7<br>±0.6 | 23<br>±2 | 34.4<br>±0.9 | 26.2<br>±1.1 | 25.00<br>±0.17 | 23<br>±3 | 16<br>±1 |

<sup>a</sup>Innis. *Adv Nutr.* 2 (2011) <sup>3</sup><sup>b</sup>Hageman et al. *Int. Dairy J.* 92 (2019) <sup>48</sup><sup>c</sup>Zhou et al. *Front. Nutr.* 11 (2024) <sup>30</sup><sup>d</sup>van Erp et al. *Metab. Eng.* 67 (2021) <sup>27</sup><sup>e</sup>Bhutada et al. *Metab. Eng. Commun.* 14 (2022) <sup>28</sup><sup>f</sup>This study<sup>g</sup>Calculated from López-López et al. *Eur. J. Clin. Nutr.* 56 (2002) <sup>102</sup><sup>h</sup>Calculated from Sun et al. *Food Chemistry* 242 (2018) <sup>103</sup><sup>i</sup>Yuhas et al. *Lipids* 41 (2006) <sup>10</sup><sup>j</sup>He et al. *Food Sci Nutr.* 12 (2021) <sup>104</sup><sup>†</sup>C10:0-C12:0
n/a = not applicable, n.d. = not determined due to minimal detection

We considered whether reducing isozyme competition from the endogenous ER-LPAAT ortholog might improve incorporation of palmitate at *sn*-2^49^. *Auxenochlorella* LPAAT2 was identified on the basis of orthology to CrLPAAT2 (Cre17.g738350)^32^, with authentic LPAAT activity in vivo (Fig S2)^47^, and the occurrence of an acyltransferase domain (IPR002123) with conserved active site motifs involved in lysophosphatidic acid substrate binding^50^. The *Auxenochlorella LPAAT2* locus is likely essential, since we could not generate *lpaat2* double knockouts unless we did so in backgrounds that also expressed the heterologous CrLPAAT2-Venus and HsAGPAT1-Venus fusions, which also validates the function of the *Auxenochlorella* ortholog (Fig. S3, S5). Endogenous *Auxenochlorella* LPAAT2-Venus and heterologous CrLPAAT2-Venus and HsAGPAT1-Venus all exhibited similar perinuclear localization, colocalizing with an mCherry ER marker (Fig. 2C and S4) at the primary site of TAG biosynthesis^51,52^. Wild type and *lpaat2* single allele knockouts exhibited similar growth (Fig. S3C) and lipid content measured as a fraction of biomass at the end of the time course (Fig. S5), but growth and lipid accumulation were reduced in *lpaat2* double knockout strains carrying the *Chlamydomonas* and human enzymes, suggesting that in single copy the heterologous *LPAAT* genes did not fully compensate for the loss of endogenous LPAAT2 function (Fig. S3C and S5). The fatty acid profiles of the modified strains were unchanged from wild type (Fig. 2D, S5). When we compared strains expressing *CrLPAAT2* or *CrLPAAT2-Venus* in the wild-type background to the strains lacking one or both endogenous *LPAAT2* alleles we noted no significant increase in C16:0 incorporation at *sn*-2 (Fig. 2D, compare JL27 to JL73; Table S3, compare JL27 to JL83 and JL83+JL33). From these results we conclude that *Auxenochlorella LPAAT2* encodes the Kennedy pathway LPAAT, and reducing competition from the endogenous isoenzyme does not substantially improve C16:0 *sn*-2 regioselectivity catalyzed by the heterologous LPAATs. CrLPAAT2 was chosen over HsAGPAT1 for further strain development because of its greater C16:0 selectivity.

### Stacking genetic modifications to mimic HMF fatty acid and regioisomeric composition

We combined the thioesterase and LPAAT activities to reconstitute the major fatty acids and TAG regioisomers of HMF in a single strain. *CwFATB2* and *BjFATB3* were co-expressed in wild type and in strains expressing *CrLPAAT2* at *dao1* (JL27) or at *lpaat2-1* (JL73) (Fig. 3A). Thioesterase-expressing strains JL52 and JL88 produced about 10−18% MCFA, and 12–22% C16:0 (Fig. 3A, Table S3), comparable to the saturated fatty acid content of HMF from Filipino mothers (Fig. 1A)^10,53^. At about 3%, C18:0 was short of the 4-6% target, likely due to competition between endogenous and heterologous thioesterase and β-ketoacyl-ACP synthase activities (Fig. 3A). C16:0 comprises 36–42% of total fatty acids at *sn*-2 in the CrLPAAT2 expressors (JL52+JL27, JL88+JL27 and JL88+JL73) (Fig. 3B, Table 1). With 12−14% MCFA, 20−21% C16:0, and 68−71% of total C16:0 at the *sn*-2 position, strains JL88+JL27 and JL88+JL73 closely approximate HMF fatty acid and *sn*-2 palmitate composition (Fig. 3A, 3B, Table 1). C16:0 incorporation at *sn*-2 was comparable in the stacked trait strain with disrupted *lpaat2-1* (JL88+JL73), compared to the equivalent strain with both *LPAAT2* alleles intact (JL88+JL27), again demonstrating that reducing competition from endogenous LPAAT2 provided no benefit (Fig. 3A, 3B, Table 1). Individually, the modifications to increase *sn*-2 palmitate (Fig. 2D, JL27), MCFA, C14:0 and C16:0 levels (Fig. 1A, JL88) had only moderate impacts on growth (Fig. 3C) and lipid titer (Fig. 3D). However, when combined in the JL88+JL27 strain, lipid titer decreased by 20–25% compared to the wild type (Fig. 3D, S6, S7). Because the lipid fraction per dry weight remained unchanged across all strains (Fig. 3E), the reduced titer in JL88+JL27 (hereafter referred to as HMF1) likely stems from slower cell division or a delayed entry into the lipogenic phase (Fig. S6C).

**Figure 3.**
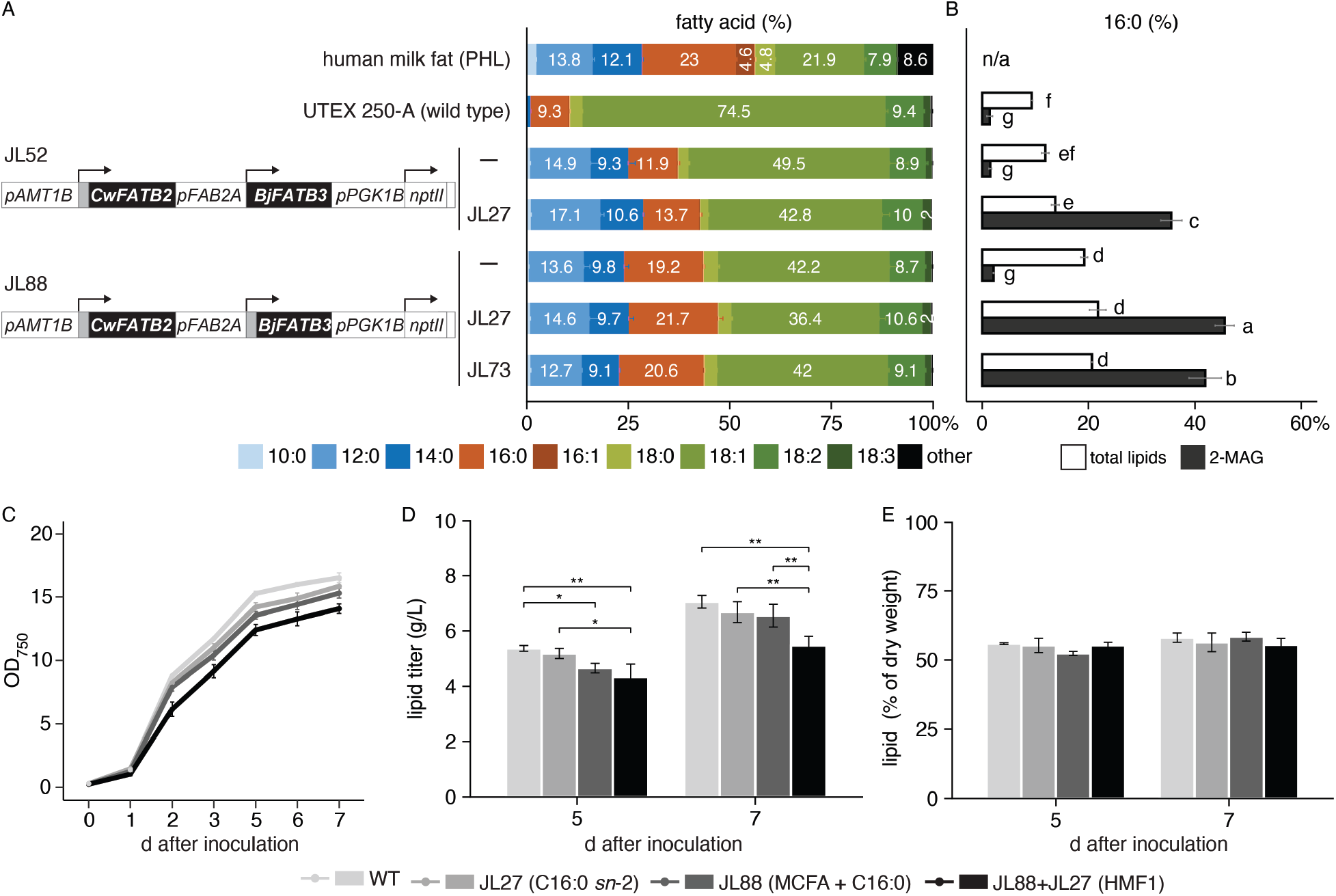
Trait stacking to mimic the medium-chain fatty acids and *sn*-2 palmitate composition of HMF. **(A)** Fatty acid composition (wt%) comparison of wild-type and engineered strains expressing thioesterases and CrLPAAT2. The thioesterase expression constructs target *CwFATB2* and *BjFATB3* to the *AMT1B-1* allele (3’ flanking DNA not shown). The *AMT1B-1* 5’ flank also serves as the promoter (*pAMT1B-1*) driving the *FAB2Atp-CwFATB2* fusion. The downstream *BjFATB3* gene encoding its native transit peptide (JL52) or fused with the FAB2A-1 transit peptide-encoding sequence (JL88) is driven by *pFAB2A-2*. The expression constructs were transformed into wild-type (—) and the *CrLPAAT2*-expressing strains JL27 and JL73. Error bars = SD. **(B)** Fraction of C16:0 in total FA (white bars) and in 2-MAG (black bars) from *sn*-2 lipase assays. Letters represent Tukey Honest Significant Differences between strains (*padj* ≥ 0.05 for measurements with the same letter). n = 3-4 independent transformants, n = 3 for independent cultures of wild type. Error bars = SD. **(C)** 7-day time course comparing growth of wild type and engineered strains after inoculation in low nitrogen (3.75 mM NH_4+_) medium. Turbidity measurements at 750 nm reflect changes in cell number and volume. Refer to Materials and Methods for detailed growth conditions. **(D)** Measurements of lipid titers from 2 mL of cell cultures in C. Error bars = SD. n = 3 independent transformations for each engineered strain and n=3 independent cultures of the wild type. Statistical significance was determined using a pairwise two-sample t-test (ns: P ≥0.05, *P < 0.05, **P < 0.01, ***P < 0.001, ****P < 0.0001). **(E)** Measurements of lipid content as a percentage of dry weight biomass from cell cultures in A. Error bars = SD. n = 3 independent transformations for transgenic strains and n=3 independent cultures of the wild type.

Linoleic acid levels are highest in HMF from Chinese and Latin American mothers, ranging from 16% to over 20% of total fatty acids^10,11,54,55^. To boost linoleic acid accumulation we introduced either *lysophosphatidylcholine acyltransferase 1* (*LPCAT1*) or *phosphatidylcholine:diacylglycerol cholinephosphotransferase 1* (*PDCT1*) from *Arabidopsis* into wild type or JL88+JL27 (Fig. 4A, 4B)^34,56,57^. Both enzymes increased linoleate abundance two to three-fold at the expense of oleic acid, but LPCAT1 activity did not interfere with C16:0 enrichment at *sn*-2, whereas PDCT1 reduced the fraction of C16:0 at *sn*-2 to 28% (Fig. 4C–E, Table 1 and S3). LPCAT1 is therefore preferred for boosting C18:2 n-6 levels in HMF strains, since it releases *sn*-2 linoleic acid from phospholipids (the site of C18:1 n-9 desaturation) into the acyl-CoA pool for subsequent incorporation into TAG (Fig. 4A)^56^. In contrast, PDCT1 catalyzes interconversion of phospholipids and diacylglycerol (DAG), and consequently competes with de novo DAG biosynthesis through the Kennedy pathway, resulting in enrichment of unsaturated fatty acids at *sn*-2 in TAGs (Fig. 4A)^58^.

**Figure 4.**
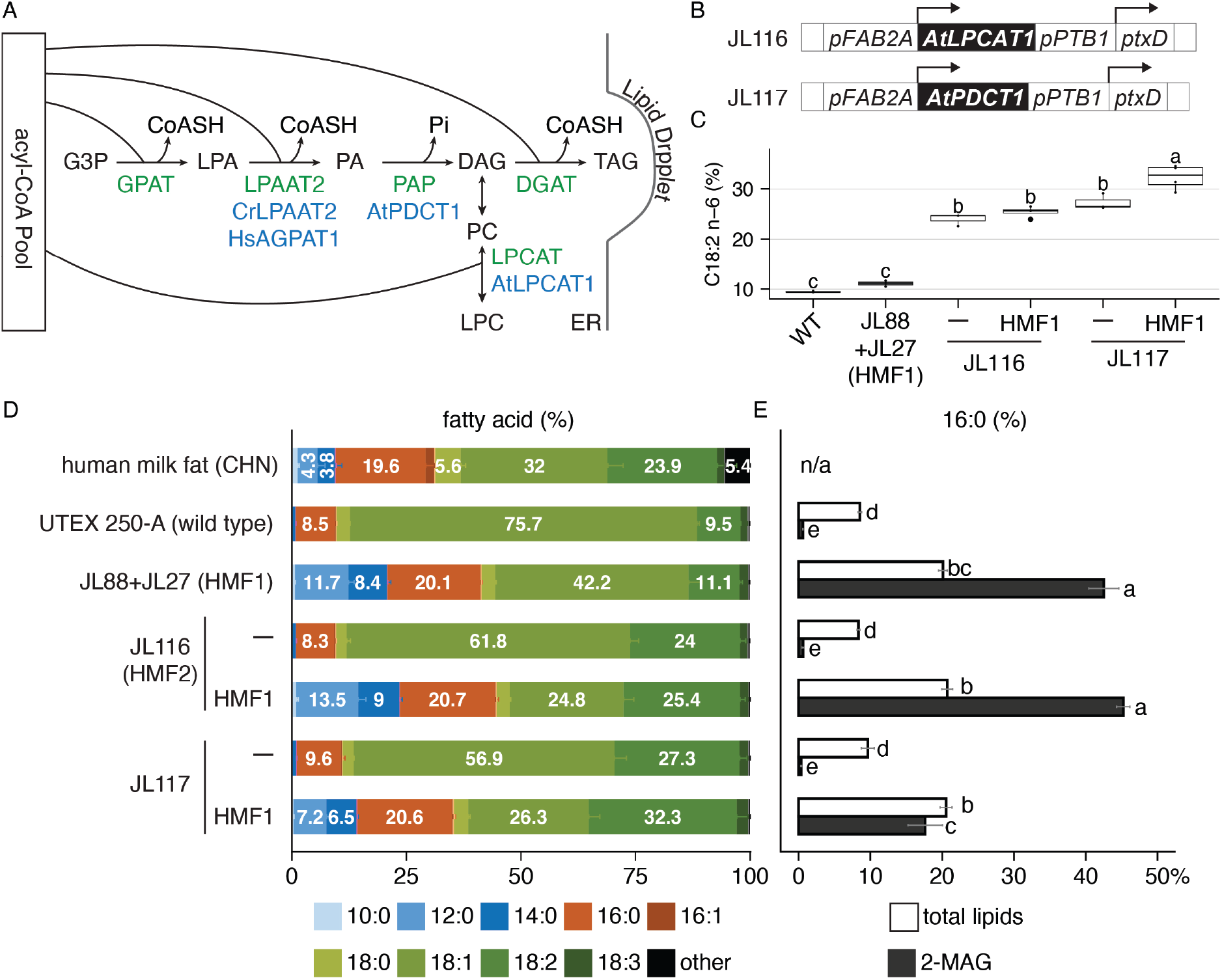
Increased incorporation of linoleic acid into TAG. **(A)** Head-group exchange between phospholipids and DAG by phosphatidylcholine diacylglycerol cholinephosphotransferase 1 (AtPDCT1) and transfer of linoleic acid from phosphatidylcholine (PC) to the acyl-CoA pool by lyso-PC acyltransferase (LPCAT1). LPC = lysophosphatidylcholine. **(B)** *AtLPCAT1* and *AtPDCT1* expression constructs. The *AtLPCAT1* (JL116) and *AtPDCT1* (JL117) constructs contain 5’ and 3’ flanking DNA targeting homologous recombination at the *THI4* locus. **(C)** C18:2 n-6 abundance (% of total fatty acids) in HMF from China (CHN), wild type UTEX 250-A and engineered strains. JL116 and JL117 were transformed into wild-type (—) and the HMF mimic strain JL88+JL27 (HMF1). **(D)** FA profile comparisons (wt%) of wild type and engineered strains. **(E)** Fraction of C16:0 in total FA (white bars) and in 2-MAG (black bars) from *sn*-2 lipase assays. n = 3-4 independent transformants of each transgenic strain. Error bars = SD. Refer to Table S1 for the strain details. n = 3 for the UTEX 250-A wild-type control. Letters represent Tukey Honest Significant Differences between strains (*padj* ≥0.05 for measurements with the same letter).

### TAG regioisomers in *Auxenochlorella* HMF strains

A targeted parallel reaction monitoring (PRM) liquid chromatography-tandem mass spectrometry (LC-MS/MS) method was developed and applied to examine the ratios of key TAG regioisomers in HMF1 and JL116+JL88+JL27 (HMF2) (Fig. 5)^59,60^. Diacylglycerol (DAG) daughter ion ratios in MS/MS spectra indicate that in wild-type lipid, 98% of dioleoyl-palmitoyl-glycerol molecules were either 1,2-dioleoyl-3-palmitoyl-glycerol (OOP), or 1-palmitoyl-2,3-dioleoyl-glycerol (POO). In contrast, in the HMF mimics OPO comprises 71% of dioleoyl-palmitoyl-glycerol TAGs (Fig. 5A). The engineered strains, designed to have an increased diversity of fatty acids, have lower levels of oleate and hence lower absolute abundance of OPO + OOP + POO (Fig. 5A). The abundance of dioleoyl-palmitoyl-glycerols is highest in the wild type because of the low complexity of the fatty acid profile (mainly oleic, palmitic and linoleic acids). Similarly, the *sn*-2 palmitate regioisomers account for 83-87% of oleoyl-dipalmitoyl-glycerols (PPO + POP + OPP) due to increased accumulation of C16:0 in engineered strains over wild type (Fig. 5B). These mass spectrometry analyses were broadly consistent with the results of orthogonal *sn*-2 lipase assays (Fig. 3B, 4E, Table S3).

**Figure 5.**
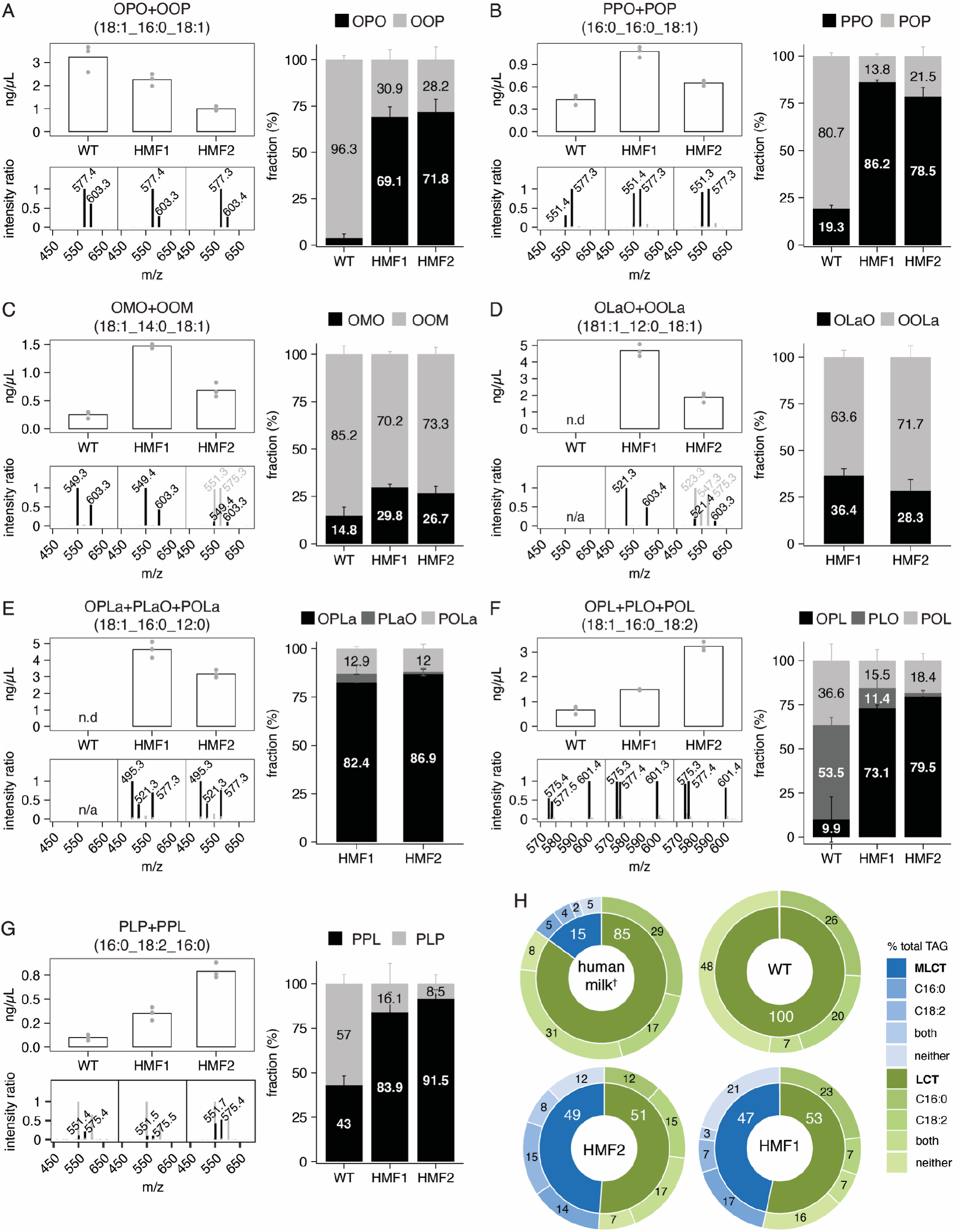
TAG regiospecificity in HMF mimic lipids. Regiospecificity of seven representative TAGs (total 16 isomers) in total lipid extracts of wild type, JL88+JL27 (HMF1), and JL116+JL88+JL27 (HMF2) were measured and quantified using targeted HPLC-MS/MS. The top-left graph in each panel presents TAG quantification based on MS1 peak height. The bottom-left graphs illustrate spectra that were used to calculate ratios of target MS2 fragments. Regioisomer ratios in the graphs on the right were calculated by comparing sample spectra with spectra from corresponding mixtures of authentic standards for each TAG species. The method does not distinguish between *sn*-1 and *sn*-3 stereoisomers, hence one stereoisomer represents both (e.g. OOP represents both OOP and POO, etc.). Error bars = SD. n = 3 independent transformants. (H) Global TAG characterization from Lipidyzer shotgun lipidomics in wild type, HMF1 and HMF2. A total of 311 TAG sub-species identified from 65 bulk species (e.g., 54:3) were quantified and categorized into MCT (all acyl chains C12), MLCT (containing both C12 and C14−20 acyl chains), and LCT (all acyl chains ≥C14). Within each group, TAGs containing at least one C16:0 and/or C18:2 were specified and further color-coded. Values represent the mean % of total TAGs from n = 4 independent transformants of each strain. Refer to Table S7 for isobaric TAG deconvolution and Supplemental dataset S1 for raw data. † Human milk data was adapted from Yu et al., 2022^64^.

When we analyzed TAGs containing C12:0 and C14:0, we found that a little over one third of lauryl-dioleoyl-glycerols and about 30% of myristoyl-dioleoyl-glycerols have the saturated fatty acid esterified at *sn*-2 (Fig. 5C, 5D, OLaO and OMO), as would be expected if CrLPAAT2 does not discriminate against C12:0 and C14:0 at that position. In contrast, 85-86% of lauryl-oleoyl-palmitoyl-glycerols contain palmitate at *sn*-2 (OPLa or LaPO) (Fig. 5E), reflecting the CrLPAAT2 preference for C16:0. Consistent with the FA profile from GC-MS, laureate-containing TAGs were not detected in wild-type lipid (Fig. 5D, 5E). The predominant regioisomers of linoleoyl-oleoyl-palmitoyl-glycerol and linoleoyl-dipalmitoyl-glycerol TAGs in the engineered lipids also contained *sn*-2 palmitate (72-76% OPL or LPO, Fig. 5F, 86-90% PPL or LPP, Fig. 5G), and as expected, increased linoleic acid in the HMF2 strain expressing LPCAT1 correlated with higher abundance of OPL (or LPO) relative to OPO, changing the OPO:OPL ratio from 1.4 in HMF1 to 0.3 in HMF2. The availability of standards limited our regiospecific PRM quantification to a few key TAG regioisomers, so to achieve a broad understanding of the abundances of TAG species in the HMF strains we undertook shotgun lipidomic analysis using the Lipidyzer platform (Fig. S9, S10)^61,62^. This method reports TAGs as mass:saturation-fatty acid (eg. TG 52:2-FA18:1, see Supplemental Dataset 1)^63^. We employed a least-square-based deconvolution method to estimate the contribution of isobaric TAGs to each of the partial molecular species (e.g. TG 40:0 = ∼75% TG 12:0_12:0_16:0 and ∼ 25% TG 12:0_14:0_14:0 in HMF1 and HMF2, Table S7). Categorization of predicted TAGs revealed that while wild type lipids were composed entirely of LCTs, about half of the TAG species in HMF1 and HMF2 were C12:0-containing MLCTs (Fig. 5H). By comparison, HMF samples from mothers in six regions in China contained an average of 16% MLCTs containing at least one C6-12:0 acyl group (Fig. 5H)^64^. Together, the targeted LC-MS/MS PRM results (Fig. 5A−G), and Lipidyzer quantification (Fig. 5H) revealed substantial increases in *sn*-2 palmitate-containing TAGs and MLCTs in engineered *Auxenochlorella*, mimicking the bulk TAG composition of HMF^65,66^.

## Discussion

We have engineered *Auxenochlorella* fatty acid and lipid biosynthesis to replicate the *sn*-2 palmitate, MCFA and linoleic acid content of HMF. By stacking genetic modifications we created a single-source algal oil that can be further tailored to match maternal geography or infant nutritional requirements at different developmental stages. Having identified the parts and approach, fully stacked HMF strains may be generated in one or two transformation steps to produce lipids that substitute for vegetable oils in formula without chemical and enzymatic processing. Increased MCFA and C16:0 in TAG were achieved through introduction of select plant thioesterases which terminate acyl chain elongation. Selectivity for palmitate esterification at *sn*-2 was realized using human or *C. reinhardtii* LPAATs. These heterologous LPAATs display ER localization patterns resembling that of the endogenous enzyme and complement the corresponding *Auxenochlorella lpaat2*-null mutant, attesting to equivalent function (Fig. 2C, D).

In strains engineered to accumulate ≥20% of C16:0, the proportion of palmitate at *sn*-2 was between 68−73%, in line with the 70−88% range of C16:0 found at *sn*-2 reported for HMF^3,48,60^. Increased palmitate incorporation at *sn*-2 to reach the upper end of the HMF range may be achieved by introducing highly selective LPAATs discovered in *Pedinophyceae* sp. or *Edaphochlamys debaryana*^29^. Alternatively, or in concert, heterologous glycerol phosphate acyltransferases (GPATs) and diacylglycerol acyltransferases (DGATs) with low preference for palmitate, such as plastidic GPATs from chilling resistant plants^67,68^, or DGATs from select plant species^69–71^ could contribute to exclusion of C16:0 from the *sn*-1 and *sn*-3 positions.

TAGs containing mixed medium- and long-chain fatty acids may be more beneficial for lipid absorption compared to trisaturated MFCA-containing TAGs from coconut or palm kernel oil ^23,66^. Indeed, up to 34% of TAG molecules in HMF contain at least one MCFA^65^. The engineered *Auxenochlorella* lipids match the relative abundance of MLCTs in HMF; notably, lauryl-dioleoyl-glycerols and lauryl-oleoyl-palmitoyl-glycerols are among the most prominent TAG species in both engineered lipid profiles (Fig. 5D, 5E, 5G, Table 1)^65^. Linoleic acid abundance in TAG was increased to match that of oleic acid through the activity of LPCAT1 (Fig. 4C, 4D), resulting in OPO:OPL ratios similar to HMF from Chinese mothers (Fig. 5A, 5F)^54^. We could not quantify TAG regioisomers containing MCFA and linoleic acid for lack of standards, but lauryl-linoleoyl-palmitoyl-glycerol and other laurate- and linoleate-containing MLCTs are significant species in HMF2 lipids (Fig. 5G, Table S7). While there is no evidence for functional differences between HMF substitutes with higher OPL versus OPO, consumers may prefer infant formulae consistent with regional variations in HMF composition.

Looking forward, a realistic objective is to provide a “whole fat” solution for infant formula by engineering algae oils that fulfill most lipid nutrition requirements, including the recommended levels of long-chain unsaturated fatty acids that are important for immune and nervous system development^2^. The endogenous ER-associated acyl-CoA elongation pathway in *Auxenochlorella* has already been co-opted and augmented with plant and microbial elongases and desaturases to produce arachidonic (C20:4n-6, ARA), eicosapentaenoic (C20:5n-3, EPA) and nervonic (C24:1n-9) acids^33,72,73^. One more two-carbon elongation and desaturation step would generate docosahexaenoic acid (C22:6n-3, DHA) from EPA^74,75^. An added benefit of *Auxenochlorella* lipids is the presence of lutein and zeaxanthin, fat-soluble xanthophylls with benefits for infant eye development and health^76^. Carotenoid supplementation would likely exceed the recommended dose if all of the fat in infant formula came from *Auxenochlorella* oil^77^, but manipulation of carotenoid biosynthesis to reduce carotenoid content and change the xanthophyll composition has been demonstrated^34^. HMF mimic oils could also be engineered to provide vitamin A through expression of β-carotene 15,15’-dioxygenase, retinaldehyde reductase, and acyltransferases for conversion to retinyl esters^78^. Collectively, the native capacity for lipid and carotenoid accumulation, coupled with tractability towards pathway engineering, position *Auxenochlorella* as a uniquely versatile production platform for lipid ingredients tailored for the complex requirements of infant formula.

## Materials and Methods

### Strains and growth conditions

All engineered strains (Table S1) were generated in the *Auxenochlorella protothecoides x symbiontica* UTEX 250-A wild-type background derived from UTEX 250 ^32^. Cells were cultivated heterotrophically in standard ApM1 medium with 2% glucose, 2 μM thiamine-HCl, and 7.5 mM (NH_4_)_2_SO_4_^79^. For lipid induction, cells were cultured in 12-well plates (Corning™ 3512) containing 1 mL of ApM1 with 0.5% glucose (other ingredients unchanged) in the dark, shaken at 150 rpm at 26°C for 3 days in an Innova 44 incubator (New Brunswick), allowing the culture to reach stationary phase. 320 μL of the stationary phase cultures were used to inoculate 4 mL ApM1, seed cultures which were grown with shaking at 150 rpm at 26^°^C for 24 h. Aliquots of the seed cultures were used to dilute cells 50-fold into a total volume of 15-25 mL of ApM1 with reduced nitrogen (3.75 mM (NH_4_)_2_SO_4_), corresponding to a quarter of the nitrogen in the standard medium to promote storage lipid accumulation (day 0). An additional 2% (w/v) glucose was added on day 2 to increase the C/N ratio and increase lipid yield. Samples were collected, unless specified, on day 5 during the lipid-accumulating phase for downstream lipid analysis.

### Auxenochlorella transformations

Transformations were performed as described by Dueñas et al.^80^. Briefly, in 1.5 mL eppendorf tubes, 15 μg (in 10-15 μL) of linearized plasmid DNA was mixed with 150 μL of cells pre-treated with 0.1 M lithium acetate/1X TE pH 8. 750 μL of 40% (v/v) polyethylene glycol-4000 in 0.1 M lithium acetate/1X TE pH 8 was then added, and the mixtures were shaken at 150 rpm in the dark at 26°C for 16 h. Cells were collected by centrifugation at 2348 x *g* for 30 s in an eppendorf 5424 tabletop centrifuge, resuspended in 0.25 mL of 1 M sorbitol, and plated on selective ApM1 agar plates^79^. Selection markers included 1) *SUC2* for growth on 2% sucrose, 2) *nptII* for selection with 100 μg/mL geneticin (G418 sulfate), 3) *THIC* for selection for growth on thiamine-free media, and 4) *ptxD* for selection for growth on phosphite^32^. All transformants were genotyped by PCR amplification and DNA sequencing of the resulting PCR products to confirm construct integration at the targeted insertion sites. At least three independent transformants from each transformation were chosen for further analysis (Table 1).

### Plasmid construction

DNA constructs were designed and managed using Geneious Prime (www.geneious.com) and SnapGene software (www.snapgene.com). All heterologous genes, listed in Table 1, were synthesized by GenScript Corporation, USA after codon-optimization to match the codon-bias in UTEX 250-A. DNA fragments were generated by PCR amplification using Herculase Fusion II DNA Polymerase (Agilent 600677) or restriction enzyme digestion of plasmid DNA, then assembled using NEBuilder® HiFi DNA Assembly Master Mix (New England Biolabs). DNA sequences encoding the predicted *FAB2A-1* (UTEX250_B12.15855.1) gene promoter (*pFAB2A*) and transit peptide (amino acids 1-38, FAB2Atp, Table 1) were amplified from UTEX 250-A genomic DNA. For the mCherry ER reporter (pJL105, Table 1), a DNA sequence encoding amino acids 1-39 of BIP1 (UTEX 250-A_A01.00955.1) was amplified from UTEX 250-A cDNA and assembled with a codon-optimized *mCherry* sequence^81^ and a 12-bp extension encoding the C-terminal HDEL retention sequence. The 1158-bp *LPAAT2* (UTEX 250-A_B12.35270.1) coding sequence was amplified from UTEX 250-A cDNA and fused to a codon-optimized *Venus* DNA sequence^81^.

### Lipid extraction and fatty acid profiling

Total lipids from lyophilized or fresh cell pellets obtained from 1-2 mL cell cultures were extracted as described by Lin et al^82^, using a modified Bligh and Dyer method, with chloroform replaced by dichloromethane^83–85^. For fatty acid (FA) composition analysis, C13:0 (Sigma Aldrich 91988, 5 mg/mL in dichloromethane) was added to the lipid extracts as an internal standard. Total FAs were derivatized by incubating in 1N methanolic hydrochloric acid at 80 °C for 30 min to generate fatty acid methyl esters (FAMEs), dissolved in hexane for GC-MS analysis (Agilent 7890A GC system connected with MSD 5977A mass spectrometer, J&W DB-FastFAME GC column 60 m, 0.25 mm, 0.25 µm). Data obtained from GC-MS were analyzed using Agilent MassHunter Workstation Software (Quantitative Analysis ver. B.07.00 and Quantitative Analysis ver. B.06.00). Peaks of individual FAs were verified by comparing their retention time (RT) with FAMEs from standard mixes (Supelco CRM47885, Cayman Chemical 29365), targeted detection of *m*/*z* in extracted ion chromatograms (EIC), and spectrum search in the NIST (National Institute of Standards and Technology Library). The resulting data (integrated peak areas) were calibrated with standard curves (peak concentration) derived from external FAME standards. Data processing, statistical analysis, and plotting were performed using R 4.2.2.

### Lipase-mediated regiochemical analysis of TAG

The complete procedure, adapted from Luddy et al., 1964^86^, is described in Lin et al^82^. Briefly, 4.5-5 mg of extracted total lipids were digested with 4 mg of *sn*-1, 3-specific porcine pancreas lipase (Sigma Aldrich L3126) in the presence of 0.018% taurocholic acid (Sigma Aldrich T4009) and 1.63% CaCl_2_ in 1 M Tris-Cl at pH 8.0 at 40 °C for 10 min. The resulting 2-monoacylglycerol (2-MAG) was separated on a thin-layer chromatography (TLC) Silica gel 60 plate (Sigma Aldrich 1057210001) in 101 mL of hexane: ethyl acetate: acetic acid = 50:50:1 (v/v) developing solvent. ^87,88^. The separated neutral lipids were visualized with 0.05% premuline (Sigma Aldrich 206865), and the 2-MAG fraction was collected from the TLC plate and extracted with dichloromethane, followed by methyl esterification for GC-MS analysis of the FA at the *sn*-2 position.

### Fluorescence microscopy of LPAAT proteins and ER structure

Engineered strains cultured under lipid-induction conditions were collected on day 5 for microscopy. Cells were embedded in 2% low-melting-point agarose and 150 mM sorbitol to minimize cell movement and to position the cells within the same focal plane on the slide. Fluorescent signals of target proteins within the cells were observed using a Zeiss LSM710 Confocal Microscope for multi-modal imaging of individual cells, utilizing single-channel confocal scanning of the individual fluorophores (mCherry: Ex/Em 561/596 nm, VENUS: Ex/Em 514/572 nm) and super-resolution imaging with the Airyscan detector. Subsequent data processing used the Zeiss ZEN Blue 3.0 software.

### Tandem mass spectrometry analysis of triacylglycerols (TAGs). Standards and samples

The following TAG isomer standards were purchased from ABITEC (Larodan, Monroe, MI): OPO (1,3-dioleoyl-2-palmitoyl glycerol; 34-1831-7), OOP (1,2-dioleoyl-3-palmitoyl glycerol; 34-1821-7), POP (1,3-palmitoyl-2-oleoyl glycerol; 34-1611-7), PPO (1,2-palmitoyl-3-oleoyl glycerol; 34-1601-7), OMO (1,3-olein-2-myristin; 34-1830-7), OOM (1,2-olein-3-myristin; 34-1820-7), OLaO (1,3-olein-2-laurin; 34-1819-7), OOLa (1,2-olein-3-laurin; 34-1817-7), OPL (1-olein-2-palmitin-3-linolein; 34-3035-7), PLO (1-palmitin-2-linolein-3-olein; 34-3010-7), POL (1-palmitin-2-olein-3-linolein; 34-3012-7), LaPO (1-laurin-2-palmitin-3-olein; 34-3039-7), POLa (1-laurin-2-olein-3-palmitin; 34-3004-7), and OLaP (1-olein-2-laurin-3-palmitin; 34-3046-7). Individual standards were first prepared as 50 ng/µL solutions in 4:1 acetone:methanol (volume/volume) and then combined into three sets of standard mixtures. The standard mixtures of Set 1 were made of solutions of 0, 1.5625, 3.125, 6.25, 12.5, and 25 ng/µL OPO, OMO, OLaO, POL, and POLa. The standard mixtures of Set 2 were pairs of OPO+OOP, POP+PPO, OMO+OOM, and OlaO+OOLa mixed in the ratios of 1:0 (100%), 1:1 (50%), and 3:1 (25%), at a total concentration of 25 ng/µL (5 mixtures for each isomer pair). Set 3 comprised individual OPL, PLO, POL, OPLa, PLaO, and POLa, each at a concentration of 25 ng/µL. For algal samples, the extracted total lipids were dissolved in 4:1 acetone:methanol to a final concentration of 50 ng/µL^89^.

### Liquid chromatography-mass spectrometry and tandem mass spectrometry

Extracted lipid samples and TAG standards were analyzed using a liquid chromatography (LC) system (1200 Series, Agilent Technologies, Santa Clara, CA) that was connected in-line with an LTQ-Orbitrap-XL mass spectrometer equipped with an electrospray ionization (ESI) source (Thermo Fisher Scientific, Waltham, MA) in the QB3 Mass Spectrometry Facility at the University of California, Berkeley. The LC system was equipped with a G1322A solvent degasser, G1311A quaternary pump, G1316A thermostatted column compartment, and G1329A autosampler unit (Agilent). The column compartment was equipped with a Restek Viva C18 column (length: 250 mm, inner diameter: 1.0 mm, particle size: 5 micrometers, part number 9514571, Restek). Ammonium formate (99%, Alfa Aesar, Ward Hill, MA), 2-propanol, methanol (Optima LC-MS grade, 99.9% minimum, Fisher, Pittsburgh, PA), and *n*-hexane (95% minimum, Thermo) were used to prepare mobile phase solvents. Mobile phase solvent A was methanol with 1 mM ammonium formate and mobile phase solvent B was 50% 2-propanol/50% *n*-hexane (volume/volume). The elution program consisted of isocratic flow at 2% (volume/volume) B for 42 min, a linear gradient to 95% B over 2 min, isocratic flow at 95% B for 4 min, a linear gradient to 2% B over 2 min, and isocratic flow at 2% B for 30 min, at a flow rate of 150 µL/min. The column compartment was maintained at 25 °C and the sample injection volume was 1 µL. External mass calibration was performed using the Pierce LTQ ESI positive ion calibration solution (catalog number 88322, Thermo). Full-scan, high-resolution mass spectra were acquired in the positive ion mode over the range of mass-to-charge ratio (*m*/*z*) = 350 to 1000, using the Orbitrap mass analyzer, in profile format, with a mass resolution setting of 60,000 (at *m*/*z* = 400, measured at full width at half-maximum peak height, FWHM). Tandem mass (MS/MS or MS2) spectra of selected TAG precursor ions, [M+NH_4_]^+^, were acquired using collision-induced dissociation (CID) in the linear ion trap, in centroid format, with the following parameters: isolation width = 3 *m*/*z* units, normalized collision energy = 30%, activation Q = 0.25, and activation time = 30 ms. Preliminary data acquisition and analysis were performed using Xcalibur software (version 2.0.7, Thermo). The raw MS data (.mzML) and the corresponding GNPS2 analysis workflow are available at: https://gnps2.org/status?task=f9656e62dafb4c56a712ae1a10b80ae3.

### LC-MS/MS Data analysis

The raw data (.RAW) were processed with MZmine 4.7.29 for mass detection and targeted feature detection, and the resulting MS2 scans (.mgf) were loaded to R with Spectra 1.20.0 for further analysis^90,91^. The stereostructure of a TAG molecule can be inferred from the relative abundance of daughter ions generated from the loss of a fatty acyl chain during fragmentation^89^. In this process, the *sn*-1 and 3 acyl groups are more susceptible to cleavage, leading to a higher ratio of *sn*-1,2 (or *sn*-2,3) to *sn*-1,3 diacylglycerols. For example, the POP precursor generates daughter ions of PO (*m*/*z* 577.5) and PP (*m*/*z* 551.5) at a ratio (PO/PP) of 4.0, while the same ratio yielded from PPO is only 1.2. Standard curves of the ratios from standard set 2 mixtures of regioisomer pairs described above were generated. The nonlinear least squares model was used to estimate the fraction of each regioisomer in the algal samples based on the measured ratio of fragments (Fig. S8 A-F). For isomers with three different acyl groups like OPL, PLO, and POL, the fraction of the three resulting fragments is determined by the fraction of each of the three isomers as follows:

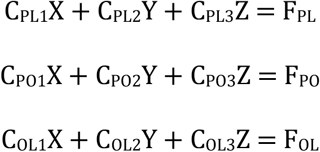

where C represents the coefficient of each fragment (PL, PO, OL), F is the resulting fraction of each fragment which is correlated to the percentage of isomer OPL (X), PLO (Y), and POL (Z). The coefficients were obtained from the fragmental fraction measured in each isomer standard (set 3), and the subsequent coefficient matrix was used for constrained non-negative least squares (NNLS) modeling to calculate the values of X, Y, and Z in algal samples (Table S5) (R package lsei 1.3)^92^.

We noticed the presence of chimeric MS2 spectra from some TAG precursors in algal samples, which was caused by co-elution of isobaric TAGs (Table S6). For example, SMO, SOM and OSM share the same precursor ion *m*/*z* with POP and PPO, which can be identified from the two additional fragment ions at *m*/*z* 549.5 (MO) and *m*/*z* 605.5 (SO) in the MS2 spectrum. However, the third fragment ion, SM, overlaps with the PP fragment ion of POP or PPO at *m*/*z* 551.5. To calculate the proportion of the fragment ion abundance at *m*/*z* 551.5 derived solely from PP, we used the following equation, assuming that the fraction of the SM fragment ion abundance (X) at *m*/*z* 551.5 corresponds to the fraction of the sum of fragment ion abundances (I) from SMO, SOM and OSM relative to the total from both SMO, SOM and OSM, and POP and PPO:

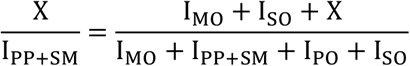

For each target TAG, we selected the four MS2 spectra exhibiting the highest cumulative calibrated peak intensities to determine average fragment ratios. TAG concentrations (ng/µL) were subsequently quantified using the highest cumulative intensity and the corresponding standard curves from standard set 1 with the four-parameter logistic model (Fig. S8G) (R package drc 3.0)^93^.

### Targeted shotgun lipidomics and data analysis

Cells harvested lipid-induction cultures described above were washed with PBS, quantified for cell density using a Nexcelom Cellometer^94^, and then stored in -80°C. Total lipids were extracted using a modified Bligh and Dyer method and analyzed on a Sciex Lipidyzer platform equipped with a Sciex 6500+ QTRAP system with differential mobility spectrometry (SelexION)^61,62^. Raw data were processed using Shotgun Lipidomics Assistant to quantitatively measure target species with internal standards and sum subclass and fatty acid content from species measurements^63^. Principal Component Analysis was performed on quantified species values with ClustVis. 311 out of 491 TAG sub-species carrying specific acyl-chains (e.g., TAG 44:0-14:0, TAG 44:0-16:0) across TG 36:0-TG 60:11 were identified in the dataset and quantified as nmol/10^7^cells^95^. For each TAG bulk species (i.e. isobaric species), we performed a deconvolutional analysis using a constrained (NNLS) algorithm^96^ to calculate the theoretical fractions of TAGs with unique acyl combinations (e.g., TAG 12:0_14:0_18:0). For example, given all the potential combinations composing a TAG 44:0 molecule, TAG 14:0_14:0_16:0, TAG 12:0_14:0_18:0, and TAG 12:0_16:0_16:0, we established the following linear equations:

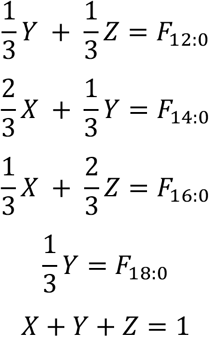

where *X*, *Y*, and *Z* represented the relative fractions of TAG 14:0_14:0_16:0, TAG 12:0_14:0_18:0, and TAG 12:0_16:0_16:0, respectively, and *F_12:0_, F_14:0_, F_16:0_, F_18:0_* denote the measured fractions of TAG 44:0-12:0, TAG 44:0-14:0, TAG 44:0-16:0, and TAG 44:0-18:0 within the TAG 44:0 bulk pool, respectively. The unknown fractions *(X*, *Y*, *Z)* were then calculated, and the model fitting was evaluated by computing the relative residual norm (E_rel_):

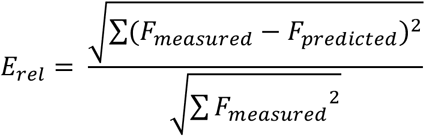

We noted that for the less abundant TAG sub-species within a bulk species (e.g., TAG 44:1-14:1 and TAG 44:1-16:1, compared to TAG 44:1-12:0), the deviation of the predicted fraction from the measured fraction was higher, which could be an artifact from the instrument detection limit. The computation was conducted using the lsei 1.3 package in R. All processed data are provided in Figure S9, S10, and Table S7.

### Sample preparation for proteomics

Cell pellets were resuspended in 8 M urea and transferred to 2 mL prefilled Micro-Organism Lysing Mix glass bead tubes. Samples were homogenized using a Bead Ruptor Elite bead mill homogenizer (OMNI International, Kennesaw, GA, USA) at speed 5.5 for 45 s. Immediately after bead beating, lysates were placed on an ice block and then centrifuged into 4 mL tubes at 1,000 × g for 10 min at 4 °C. Protein concentrations were determined using a bicinchoninic acid (BCA) assay according to the manufacturer’s instructions (Thermo Scientific, Waltham, MA, USA). For protein reduction, dithiothreitol (DTT) was added to a final concentration of 10 mM, and samples were incubated at 60 °C for 30 min with constant shaking at 800 rpm. Iodoacetamide (IAA) was then added to a final concentration of 40 mM, and samples were incubated for 45 min at room temperature in the dark. Samples were diluted to 4 M urea with 100 mM NH₄HCO₃ and 1 mM CaCl₂, followed by digestion with Lys-C (Fujifilm Wako, Richmond, VA, USA) at a 1:20 enzyme-to-protein ratio (w/w) for 3 h at 37 °C. Samples were then further diluted to 1 M urea and digested with sequencing-grade modified trypsin (Promega, Madison, WI, USA) at a 1:50 enzyme-to-protein ratio (w/w) for 12 h at 37 °C with 800 rpm shaking. Digested peptides were desalted using a 4-probe positive-pressure Gilson GX-274 ASPEC™ system (Gilson Inc., Middleton, WI, USA) equipped with Discovery C18 50 mg/1 mL solid-phase extraction tubes (Supelco, St. Louis, MO, USA). Columns were conditioned with 3 mL methanol and equilibrated with 3 mL 0.1% trifluoroacetic acid (TFA) in H₂O. Samples were loaded onto the columns and washed with 4 mL 95:5 H₂O:acetonitrile (ACN) containing 0.1% TFA. Peptides were eluted with 1 mL 80:20 ACN:H₂O containing 0.1% TFA. Eluted peptides were concentrated to approximately 100 µL using a SpeedVac concentrator. Peptide concentrations were determined by BCA assay, and samples were diluted to 0.10 µg/µL with nanopure water prior to proteomics LC-MS/MS analysis.

### LC-MS/MS of peptides

A Waters nano-Acquity dual pumping Ultra Performance Liquid Chromatography system (Milford, MA) was configured for on-line trapping of a 5 µL injection at 5 µL/min for 10 min followed by reverse-flow elution onto the analytical column at 200 nL/min. The trapping column was slurry packed in-house using 360 µm outside diameter (o.d.) x 150 µm inside diameter (i.d.) fused silica (Polymicro Technologies Inc., Phoenix, AZ) Jupiter 5µm octadecylsilane C18 media (Phenomenex, Torrence, CA) with 2 mm sol-gel frits on either end. The analytical column was slurry packed in-house using Waters BEH 1.7 µm particles packed into a 35 cm long, 360 µm o.d. x 75 µm i.d. column with an integrated emitter (New Objective, Inc., Littleton, MA). Mobile phases consisted of (A) 0.1% formic acid in water and (B) 0.1% formic acid in acetonitrile with the following gradient profile (min, %B): 0, 1; 10, 8; 105, 25; 115, 35; 120, 75; 123, 95; 129, 95; 130, 50; 132, 95; 138, 95; 140,1. MS analysis was performed using an Thermo Eclipse mass spectrometer (Thermo Scientific, San Jose, CA). The ion transfer tube temperature and spray voltage were 300°C and 2.4 kV, respectively. Data were collected for 180 min following a 27 min delay from when the trapping column was switched in line with the analytical column. High-Field Asymmetric Waveform Ion Mobility Spectrometry (FAIMS) was used at compensation voltages -40V, -60V, and -80V. Fourier Transform Mass Spectrometry (FT-MS) spectra were acquired from 300-1800 m/z at a resolution of 120k (automatic gain control (AGC) target 4 x 10^5^) and while the top 12 Fourier Transform Higher-energy Collisional Dissociation (FT-HCD)-MS/MS spectra were acquired in data dependent mode with an isolation window of 0.7 m/z and at an orbitrap resolution of 50K (AGC target 5 x 10^4^) using a fixed collision energy (HCD) of 32% and a 30 s exclusion time.

### Database Searching and Sequence Identification

Raw LC-MS/MS files were searched using the MSGFPlus algorithm ^97^ against the *Auxenochlorella* UTEX 250-A v1.1 database (16,930 entries) ^32^ supplemented with a library of commonly observed contaminant proteins (Trypsin, Keratins, etc; 16 entries). Search parameters included a parent mass tolerance of ±20 ppm and dynamic oxidation of methionine as a variable modification. A target/decoy approach was employed to assess the False Discovery Rate (FDR). Precursor ion abundances were extracted from Selected Ion Chromatograms (SICs) using MASIC software, specifically calculating the “StatMomentsArea” (area under the elution curve) ^98^. Data were collated using MAGE Extractor and File Processor (https://github.com/PNNL-Comp-Mass-Spec/Mage) and managed within a centralized SQL Server database. Resulting Peptide-to-Spectrum Matches (PSMs) were filtered to a 1% FDR (Q-value ≤ 0.01029 yielding 8.923 decoy PSMs out of 892,326 total filter passing) to generate a high-confidence list of peptide identifications per sample.

### Protein Inference and Parsimony Mapping

Clean peptide sequences (pre- and suffix amino acids as well as modification callouts) were mapped back to the search FASTA using Protein Coverage Summarizer. To resolve the degeneracy of peptide-to-protein relationships, a hierarchical parsimony approach was implemented to derive a “BestGeneID” for each sequence: 1) Unique Mapping: If a sequence mapped to only one GeneID, it was assigned as the BestGeneID. 2) Unique Sequence Preference: If multiple GeneIDs were associated, but only one possessed a unique clean sequence elsewhere, that GeneID was selected. 3) Frequency of Observation: If neither condition was met, the GeneID with the highest count of other associated clean sequences was chosen. 4) Database Index: In cases of remaining ties, the GeneID appearing first in the indexed FASTA file was assigned. Highly non-unique sequences (mapping to multiple GeneIDs with unique sequences) were excluded from further quantitation (0.61% of data).

### Data Normalization and Statistical Assessment

Bioinformatic processing was performed using Inferno (a .NET wrapper for R) ^99^. Summed peptide abundances were log_2_ transformed and normalized using a mean central tendency approach. Global data quality was assessed via box-plots, Pearson correlation heatmaps, and 2D/3D Principal Component Analysis (PCA). For protein-level quantification, normalized peptide abundances were de-logged, summed according to their assigned BestGeneID, and re-transformed to log_2_ values. Differential expression was assessed via pairwise comparisons of biological replicates (n=4). Proteins required observation in at least two replicates to pass the averaging step. A Limit of Quantitation (LOQ) was defined as two standard deviations below the mean of all averaged abundances. Statistical significance was determined using a log_2_ ratio distribution-based T-test (Z-score). GeneIDs were considered significantly differentially expressed if they met the following criteria: a |Z-score| > 2, a corresponding *p*-value < 0.05, and identification based on more than one unique peptide. Results were categorized into 18 assessment tiers to facilitate comparative analysis across all experimental conditions.

## Acknowledgements

JYTL was supported, in part, by a Postdoctoral Fellowship award (5F32HD114387) from the National Institutes of Health (NIH). MAD was supported, in part, by the Genetics Immersion Training Fellowship (1T32GM132022-01) from the NIH. The proteomics work on thioesterases was supported in part by a Department of Energy (DOE) Office of Science Biological and Environmental Research grant DE-SC0023027 to JLM, SSM, TRN. We thank Mary Lipton of Pacific Northwest National Laboratory who facilitated the proteomic analyses by SOP and CDN with support from DE-SC0023027, performed at the Environmental Molecular Sciences Laboratory, a DOE Office of Science User Facility sponsored by the Biological and Environmental Research program under Contract No. DE-AC05-76RL01830. With support from DE-SC0023027, lipidomics at Lawrence Berkeley National Laboratory (LBNL) by TRN and SMK was performed under Contract No. DE-AC02-05CH11231 with the U.S. DOE. The QB3 Mass Spectrometry Facility received NIH support (Award Number 1S10OD020062-01). The confocal fluorescence microscopy work at the Molecular Foundry was supported by the Office of Science, Office of Basic Energy Sciences, of the U.S. DOE under Contract No. DE-AC02-05CH11231, under user proposal numbers MFP-9130 and MFP-10111 supervised by Crysten Blaby. We thank Behzad Rad for technical support with confocal fluorescence microscopy. We thank the department of Chemical and Biomolecular Engineering in the College of Chemistry at the University of California, Berkeley for supporting KK through the Master of Bioprocess Engineering program, and Frederick Twigg for technical assistance with the bioreactor experiments. Shotgun lipidomic profiling was performed by the University of California, Los Angeles - UCLA Lipidomics Lab, RRID:SCR_019210.

