## Supplemental Figures for "Bioengineered algal lipids enriched in structured medium- and long-chain triacylglycerols, linoleate, and *sn*-2 palmitate for human milkfat substitutes"

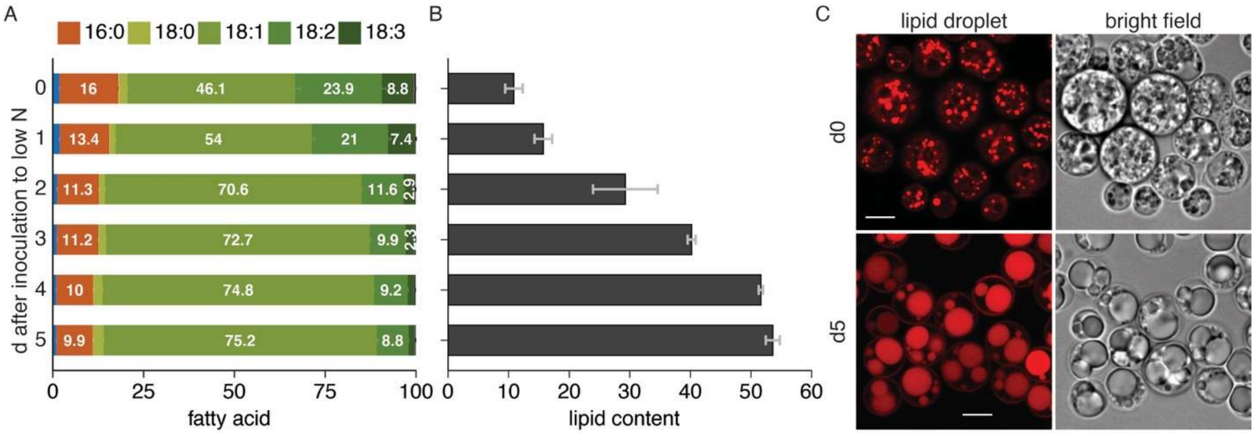


**Figure S1. Lipid accumulation in *Auxenochlorella* cells.** Changes in **(A)** fatty acid composition (wt% of total FA), **(B)** lipid content (% per dry weight), and **(C)** Lipid droplet morphology over a 5-day growth period of heterotrophic cell culture after transfer to low-nitrogen (3.75 mM $NH_4^+$) medium. Refer to Materials and Methods for detailed growth conditions of cultures and microscopy. Staining cells on day 0 and 5 with LipidTOX$^{TM}$ and observed using confocal microscopy. Scale bar = 5 µm.

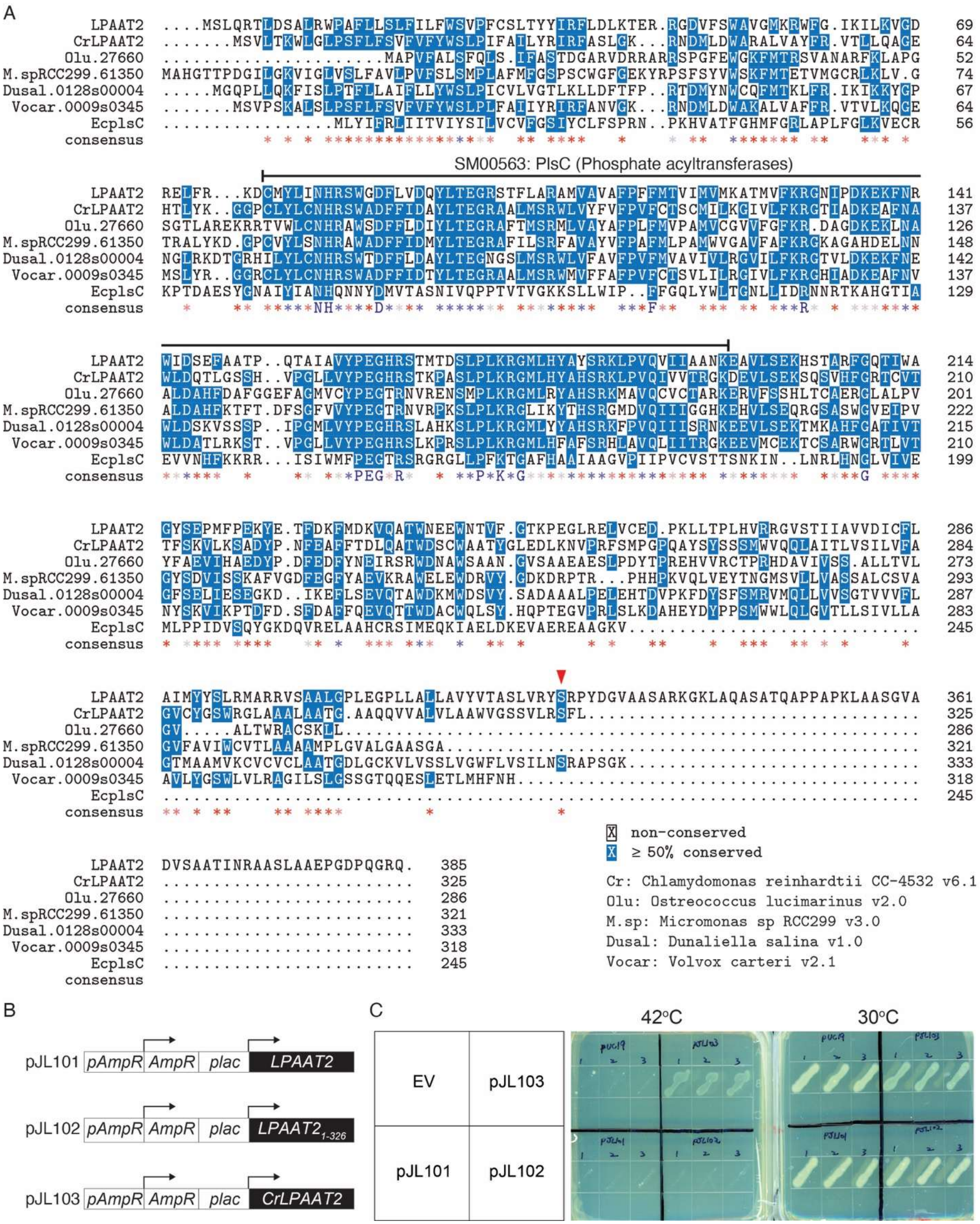
A
LPAAT2 ....MSLQRTLDSALRWPAFLLSLFILFWSVPFCSLTYYIRFLDLKTER.RGDVFSWAVGMKRWFG.IKILKVGD 69
CrLPAAT2 .......MSVLTKWLGLPSFLFSVFVFYWSLPIFAILYRIRFASLGK...RNDMLDWARALVAYFR.VTLLQAGE 64
Olu.27660 ....MAPVFALSFQLS.IFASTDGARVDRRARRSPGFEWGKFMTRSVANARFKLAPG 52
M.spRCC299.61350 MAHGTTPDGILGKVIGLVSLFAVLPVFSLSMPLAFMFGSPSCWGFGEKYRPSFSYVWSKFMTETVMGCRLKLV.G 74
Dusal.0128s00004 .....MGQPLLQKFISLPTFLLAIFLLYWSLPICVLVGTLKLLDFTFP..RTDMYNWCQFMTKLFR.IKIKKYGP 67
Vocar.0009s0345 .......MSVPSKALSLPSFLFSVFVFYWSLPLFAIIYRIRFANVGK...RNDMLDWAKALVAFFR.VTVLKQGE 64
EcplsC ....MLYIFRLIITVIYSILVCVFGSIYCLFSPRN..PKHVATFGHMFGRLAPLFGLKVECR 56
consensus
SM00563: PlsC (Phosphate acyltransferases)
LPAAT2 RELFR...KDCMYLINHRSWGDFLVDQYLTEGRSTFLARAMVAVAFPFFMTVIMVMKATMVFKRGNIPDKEKFNR 141
CrLPAAT2 HTLYK..GGPCLYLCNHRSWADFFIDAYLTEGRAALMSRWLVYFVFPVFCTSCMILKGIVLFKRGTIADKEAFNA 137
Olu.27660 SGTLAREKRRTVWLCNHRAWSDFFLDIYLTEGRAFTMSRMLVAYAFPLFMVPAMVCGVVFGFKR.DAGDKEKLNA 126
M.spRCC299.61350 TRALYKD.GPCVYLSNHRAWADFFIDMYLTEGRAFILSRFAVAYVFPAFMLPAMWVGAVFAFKRGKAGAHDELNN 148
Dusal.0128s00004 NGLRKDTGRHILYLCNHRSWTDFFLDAYLTEGRNGSLMSRWLFAVFPVFVMVAVIVLRGVILFKRGTVLDKEKFNE 142
Vocar.0009s0345 MSLYR..GGRCLYLCNHRSWADFFIDTYLTEGRAALMSRWMVFFAFPVFCTSVLILRGIVLFKRGHIADKEAFNV 137
EcplsC KPTDAESYGNAIYIANHQNNYDMVTASNIVQPPTVTVGKKSLLWIP..FFGQLYWLTGNLLIDRNNRTKAHGTIA 129
consensus NH D F R
LPAAT2 WIDSEFAATP..QTAIAVVYPEGHRSTMTDSLPLKRGMLHYAYSRKLPVQVIIAANKEAVLSEKHSTARFGQTIWA 214
CrLPAAT2 WLDQTLGSSH..VPGLLVYPEGHRSTKPASLPLKRGMLHYAHSRKLPVQIVVTRGKDEVLSEKSQSVHFGRTCVT 210
Olu.27660 ALDAHFDAFGGEFAGMVCYPEGTRNVRENSMPLKRGMLRYAHSRKMAVQCVCTARKERVFSSHLTCAERGLALPV 201
M.spRCC299.61350 ALDAHFKTFT.DFSGFVVYPEGTRNVRPKSLPLKRGLIKYTHSRGMDVQIIIGGHKEHVLSEQRGSASWGVEIPV 222
Dusal.0128s00004 WLDSKVSSSP..IPGMLVYPEGHRSLAHKSLPLKRGMLYYAHSRKFPVQIIISRNKEEVLSEKTMKAHFGATIVT 215
Vocar.0009s0345 WLDATLRKST..VPGLLVYPEGHRSLKPRSLPLKRGMLHFAFSRHLAVQLIITRGKEEVMCEKTCSARWGRTLVT 210
EcplsC EVVNHFKKRR...ISIWMFPEGTRSRGRGLLPFKTGAFHAAIAAGVPIIPVCVSTTSNKIN..LNRLHNGLVIVE 199
consensus PEG R P K G G
LPAAT2 GYSEPMFPEKYE.TFDKFMDKVQATWNEEWNTVF.GTKPEGLRELVCED.PKLLTPLHVRRGVSTIIAVVDICFL 286
CrLPAAT2 TFSKVLKSADYP.NFEAFFTDLQATWDSCWAATYGLEDLKNVPRFSMPGPQAYSYSSMWVQQLAITLVSILVFA 284
Olu.27660 YFAEVIHAEDYP.DFEDFYNEIRSRWDNAWSAAN.GVSAAEAESLPDYTPREHVVRCTPRHDAVIVSS.ALLTVL 273
M.spRCC299.61350 GYSDVISSKAFVGDFEGFYAEVKRAWELEWDRVY.GDKDRPTR...PHHPKVQLVEYTNGMSVLLVASSALCSVA 293
Dusal.0128s00004 GFSELIESEGKD..IKEFLSEVQTAWDKMWDSVY.SADAAALPELEHTDVPKFDYSFSMRVMQLLVVSGTVVVFL 287
Vocar.0009s0345 NYSKVIKPTDFD.SFDAFFQEVQTTWDACWQLSY.HQPTEGVPRLSLKDAHEYDYPPSMWWLQLGVTLLSIVLLA 283
EcplsC MLPPIDVSQYGKDQVRELAAHCRSIMEQKIAELDKEVAEREAAGKV.... 245
consensus
LPAAT2 AIMYYSLRMARRVSAALGPLEGPLLALLAVYVTASLVRYSRPYDGVAASARKGKLAQASATQAPPAPKLAASGVA 361
CrLPAAT2 GVCYGSWRGLAAALAATG.AAQQVVALVLAAWVGSSVLRSFL.... 325
Olu.27660 GV.....ALTWRACSKLL.... 286
M.spRCC299.61350 GVFAVIWCVTLAAAAMPLGVALGAASGA.... 321
Dusal.0128s00004 GTMAAMVKCVCVCLAATGDLGCKVLVSSLVGWFLVSILNSRAPSGK.... 333
Vocar.0009s0345 AVLYGSWLVLVRAGILSLGSSGTQQESLETLMHFNH.... 318
EcplsC .... 245
consensus
X non-conserved
X ≥ 50% conserved
LPAAT2 DVSAATINRAASLAAEPGDPQGRQ. 385
CrLPAAT2 .... 325
Olu.27660 .... 286
M.spRCC299.61350 .... 321
Dusal.0128s00004 .... 333
Vocar.0009s0345 .... 318
EcplsC .... 245
consensus
Cr: Chlamydomonas reinhardtii CC-4532 v6.1
Olu: Ostreococcus lucimarinus v2.0
M.sp: Micromonas sp RCC299 v3.0
Dusal: Dunaliella salina v1.0
Vocar: Volvox carteri v2.1
B
pJL101 pAmpR AmpR plac LPAAT2
pJL102 pAmpR AmpR plac LPAAT2 1-326
pJL103 pAmpR AmpR plac CrLPAAT2
C
42°C
30°C
EV
pJL103
pJL101
pJL102

**Figure S2. Sequence and functional characterization of LPAAT2 in UTEX 250-A.**

**(A)** Multiple sequence alignment of the putative LPAAT2 in UTEX 250-A and LPAAT/AGPAT orthologs from other Chlorophyta and PlsC from *E. coli*. The predicted functional domain SM00563 (PlsC) / IPR002123 (Phospholipid/glycerol acyltransferase) of LPAAT2 is indicated. Conserved residues are marked with '＊', which is scaled from red (low) to blue (high) based on the conservation score. Consensus residues are marked with uppercase letters. Red triangle indicates the S326 residue of LPAAT2. The peptide sequences were obtained from Phytozome 14 (Chlorophyte orthologs) and UniProt (EcPlsC), and the sequence alignment was performed using the CLUSTAL 2.1 algorithm with the msa 1.42 package [12]. **(B)** LPAAT2 and CrLPAAT2 expression constructs for complementation test of the *E. coli plsc* mutant. The sequences have been codon-optimized for expression in *E. coli.* pJL102 contains the truncated LPAAT2 with deletion of the 327-385 region. **(C)** The expression vectors, including the empty vector (EV, pUC19), were transformed into the *E. coli* plsc mutant (strain ID: SMC2-1) and grown at 30$^{o}$C or 42$^{o}$C.

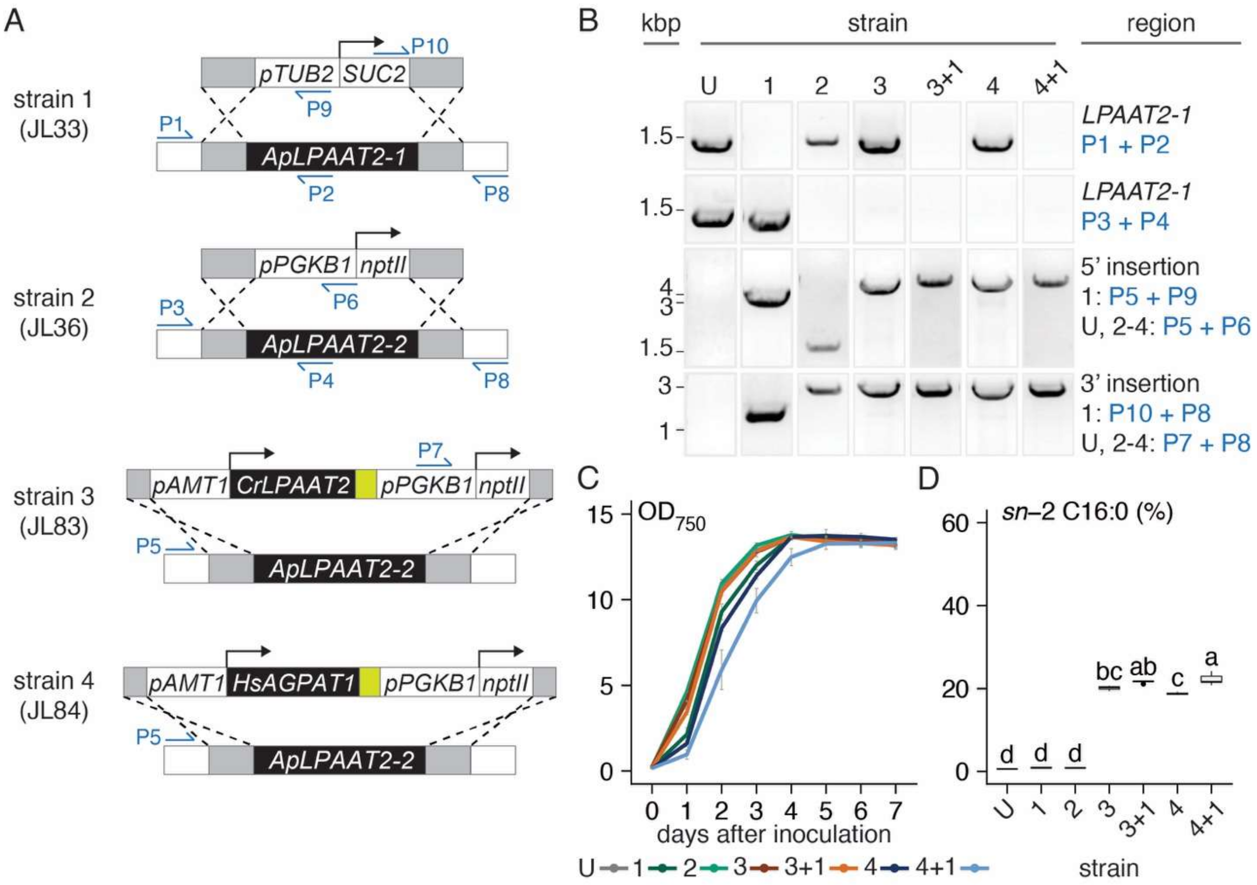


**Figure S3. Heterologous expression of LPAAT2 in *lpaat2* mutants of UTEX 250-A.** **(A)** Schematic of the constructs and landing sites. Yellow bricks represent *Venus.* Primers designed for PCR-genotyping the resulting strains are indicated by red arrows (Table S3). The endogenous *LPAAT2* alleles 1 and 2 are disrupted in *lpaat2-1*(JL33; strain 1) and *lpaat2-2* (JL36; strain 2), respectively. The CrLPAAT2-Venus construct (JL83) is introduced into the *LPAAT2-2* locus in UTEX 250-A and *lpaat2-1* (strain 1) to generate *CrLPAAT2-Venus::lpaat2-2* (strains 3) and *CrLPAAT2-Venus::lpaat2* (3+1), respectively. **(B)** PCR-genotyping of strains 1-3 to verify the disruptions in endogenous *LPAAT2* and *CrLPAAT*. **(C)** Growth curves of strain 1-4 and UTEX 250-A under low-N conditions over a 7-day course. Day 0 = the day of inoculation to the low-N medium. **(D)** 16:0 content (% of total FA) measured from 2-MAG. Letters represent Tukey Honest

Significant Differences between strains (*padj* ≥0.05 for measurements with the same letter). n = 3-5. Refer to Table S1 (strain list) and S3 (compiled data) for details.

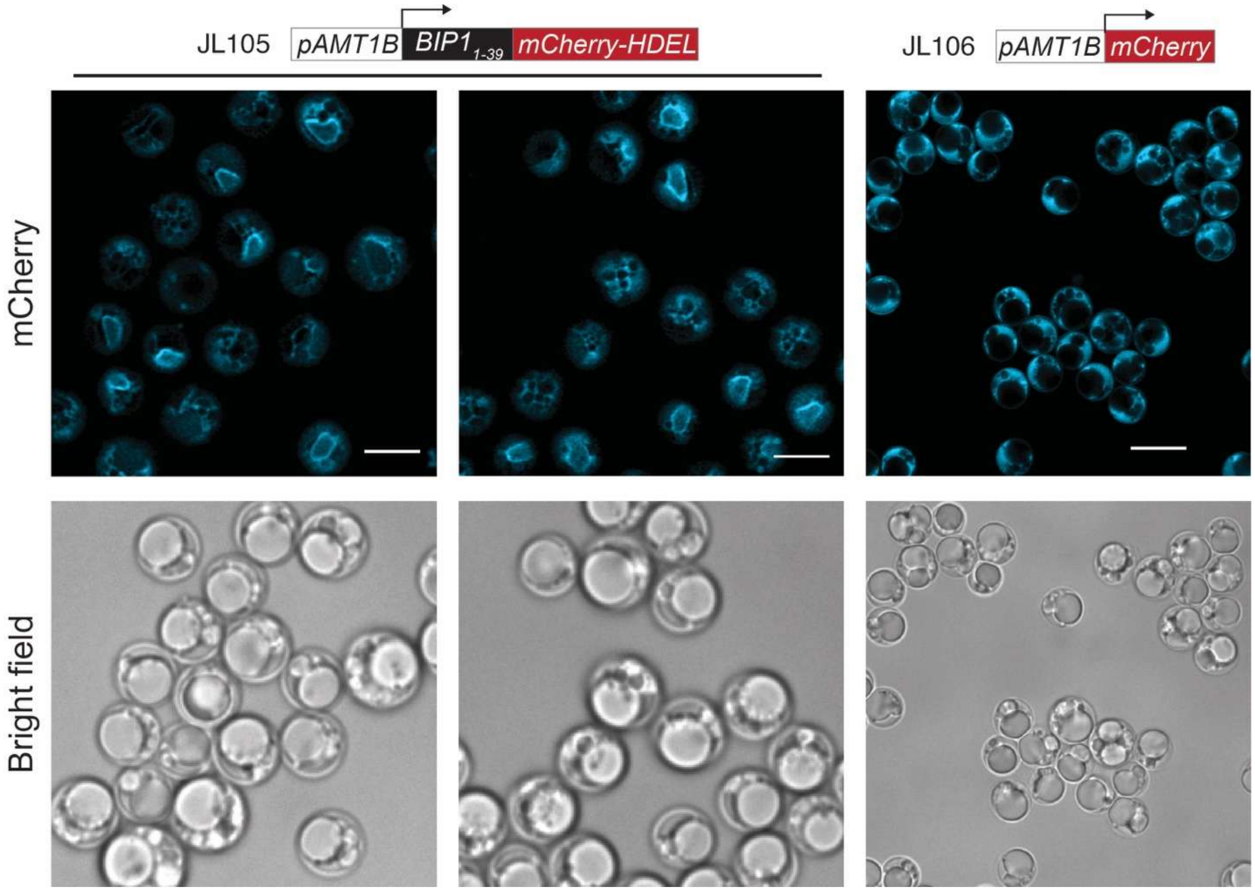


**Figure S4. ER-localization of mCherry fluorescent reporter.** Fluorescence confocal microscopy of BIP1 ER-signal peptide-tagged mCherry-HDEL marker (JL105) and mCherry-only control (JL106). Scale bar = 5 μm. Cells were grown in the lipid production phase for 5 days.

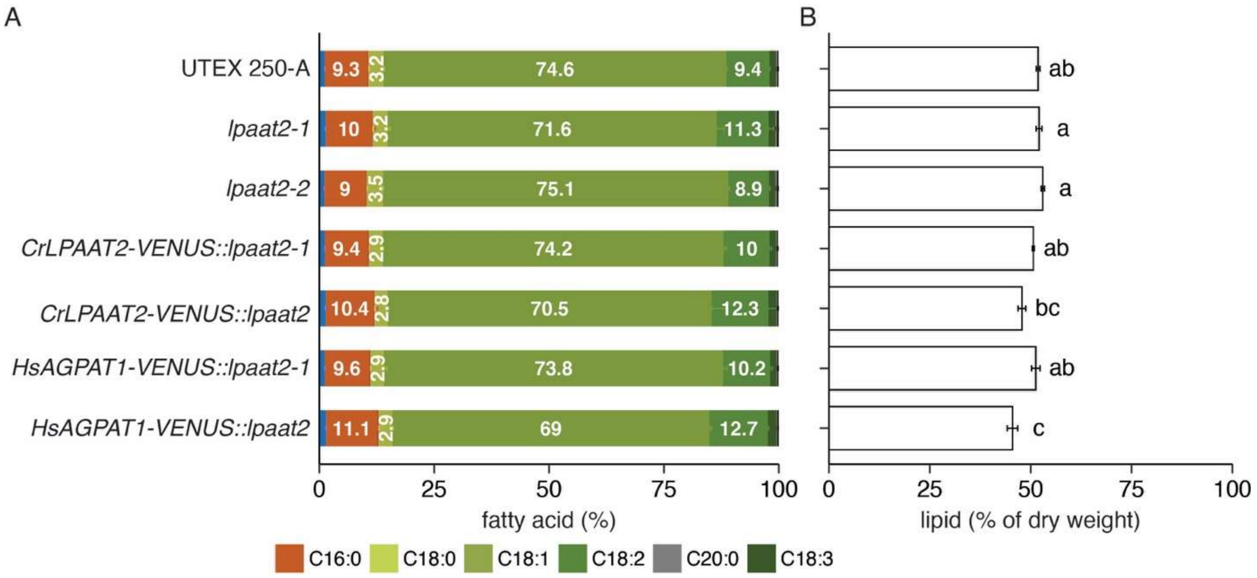


**Figure S5. Lipid accumulation in *lpaat2* complementation strains. (A)** fatty acid composition (wt%) and **(B)** lipid content (% per dry weight) of *CrLPAAT-Venus* and *HsAGPAT1*-expressing strains and their respective controls from day-5 cell cultures in Fig. 5. Letters represent Tukey Honest Significant Differences between strains (*padj* ≥0.05 for measurements with the same letter). n = 3-4. Refer to Table S1. for the strain details.

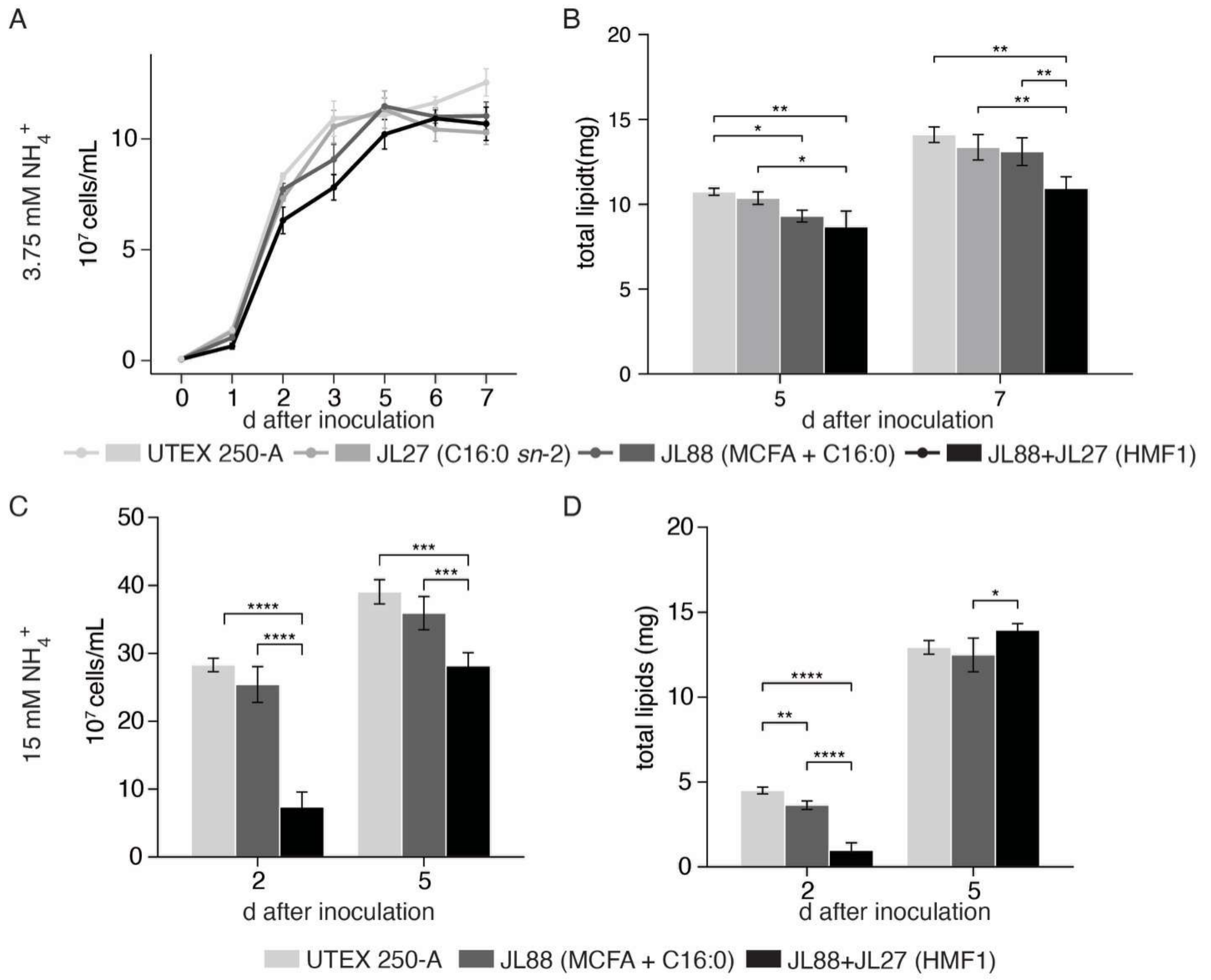


**Figure S6. Cell growth and lipid yield of HMF-producing UTEX 250-A strains. (A)** Cell density measurements of the wild type (UTEX 250-A) and engineered strains of *CrLPAAT2* (JL27), *CwFATB2* + *BjFATB3* in the wild-type (JL88) and JL27 (JL88+JL27) backgrounds in low nitrogen medium. **(B)** Measurements of total lipids (mg) from 2 mL of cell cultures in A. Error bars = SD. n = 3 independent transformations for each engineered strain and n=3 independent cultures of the wild type. **(C)** Cell density measurements of the UTEX 250-A and transgenic strains of *CwFATB2* + *BjFATB3* in the wild-type (JL88) and JL27 (JL88+JL27) backgrounds in nitrogen-replete medium. **(D)** Measurements of total lipid yield (mg) extracted from 2 mL of cell cultures shown in part C. Error bars = SD. n = 6 (JL88) and 4 (JL88+JL27) independent transformations for engineered strains and n=3 independent cultures of the wild type. Statistical significance was determined using a pairwise two-sample t-test (ns: $P \geq 0.05$, $^{*}P < 0.05$, $^{**}P < 0.01$, $^{***}P < 0.001$, $^{****}P < 0.0001$).

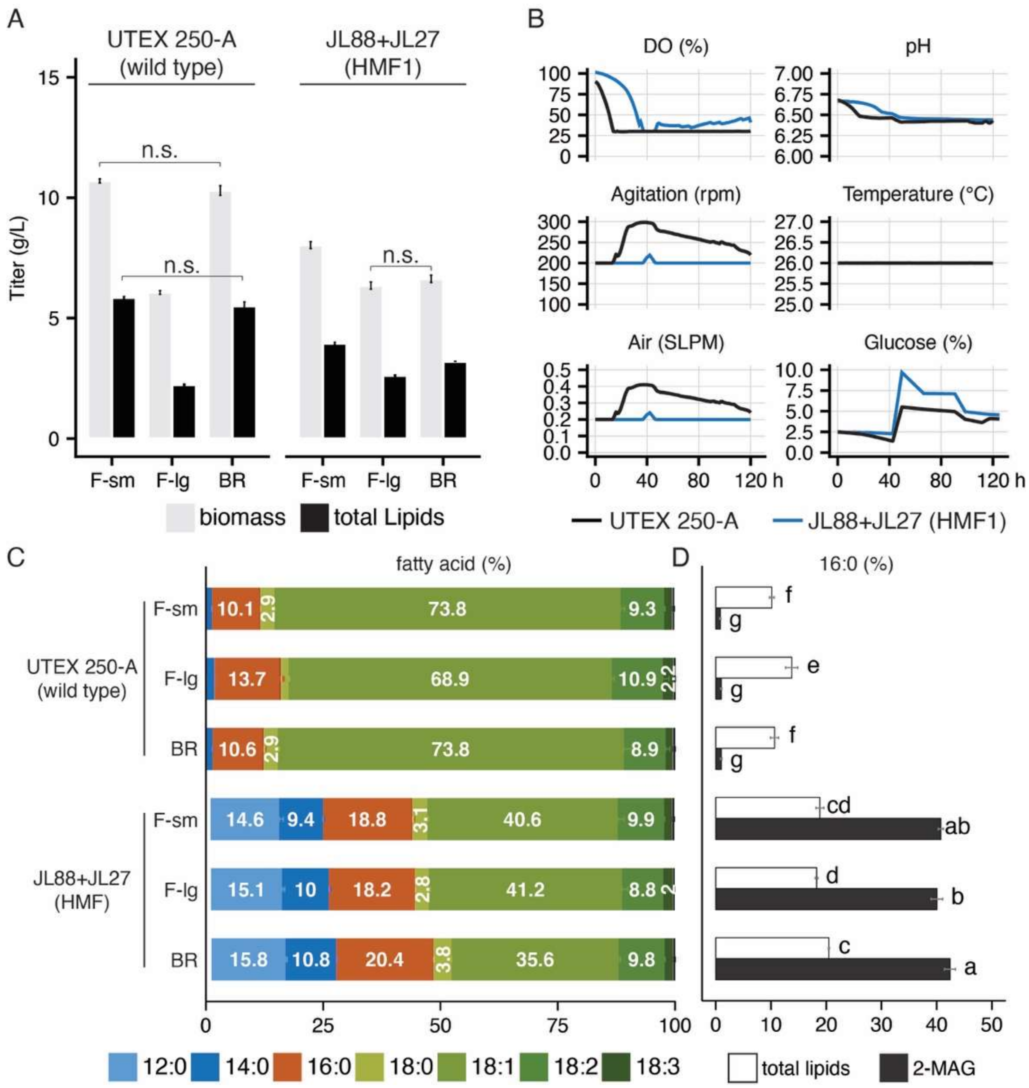


**Figure S7. Scaling up cultivation of HMF-engineered strain for lipid production.** The engineered strain JL88+JL27 and wild-type control UTEX 250-A were inoculated in low-N ApM1 medium at identical inoculation ratios (v/v) in three vessel types: small flask (F-sm, 25 mL culture volume, 125 mL capacity), large flask (F-lg, 2 L culture volume, 2.8 L capacity), and bioreactor (BR, 2 L culture volume, 3 L capacity), as described in Materials and Methods. **(A)** Biomass and lipid titers (g/L) measurements of the day-5 cell cultures of different vesicle types **(B)** Bioreactor data recorded over a 5d (120 h) course in the low-N medium. DO%: dissolved $O_2$, Air (Standard Litres Per Minute or SLPM): 2.5% (w/v)

glucose was provided on d 0 and gradually increased to 4% (UTEX 250-A) and 9.7% (JL88+JL27). **(C and D)** FA composition (wt%) and 16:0 content measured from total lipids (white) and 2-MAG (black).

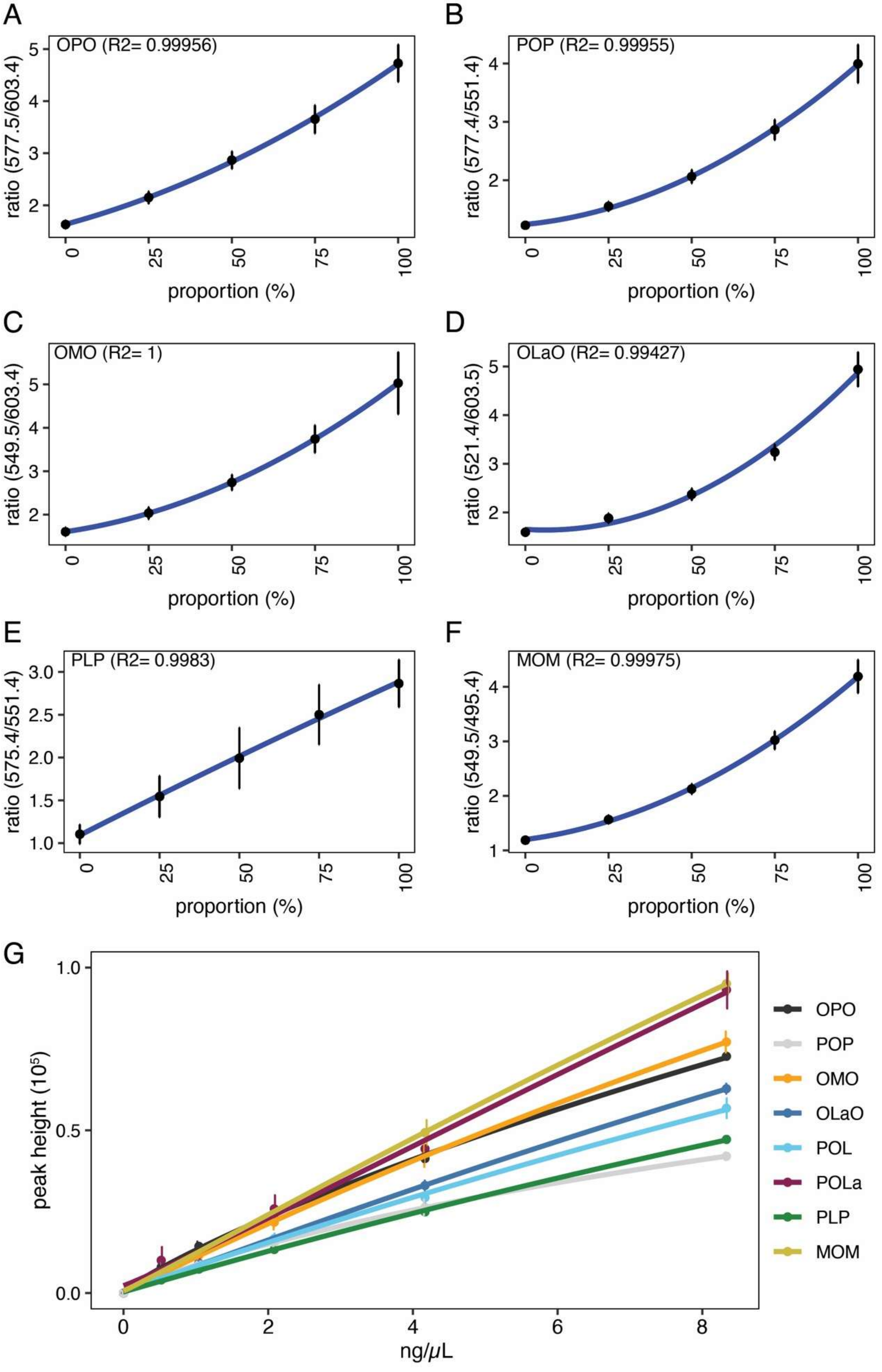
A
OPO (R2= 0.99956)
ratio (577.5/603.4)
proportion (%)
B
POP (R2= 0.99955)
ratio (577.4/551.4)
proportion (%)
C
OMO (R2= 1)
ratio (549.5/603.4)
proportion (%)
D
OLaO (R2= 0.99427)
ratio (521.4/603.5)
proportion (%)
E
PLP (R2= 0.9983)
ratio (575.4/551.4)
proportion (%)
F
MOM (R2= 0.99975)
ratio (549.5/495.4)
proportion (%)
G
peak height (10^5)
ng/μL
OPO
POP
OMO
OLaO
POL
POLa
PLP
MOM

**Figure S8. Standard curves for quantification of TAG regioisomers.** (A-F) Non-linear least squares regression of calibrated daughter-ion intensity ratios from target TAG regioisomer mixtures. x-axis: fraction of the title-indicated regioisomer in the mixture, e.g., panel A = OPO. (G) Four-parameter logistic curves of cumulative calibrated daughter-ion intensities against target TAG concentration.

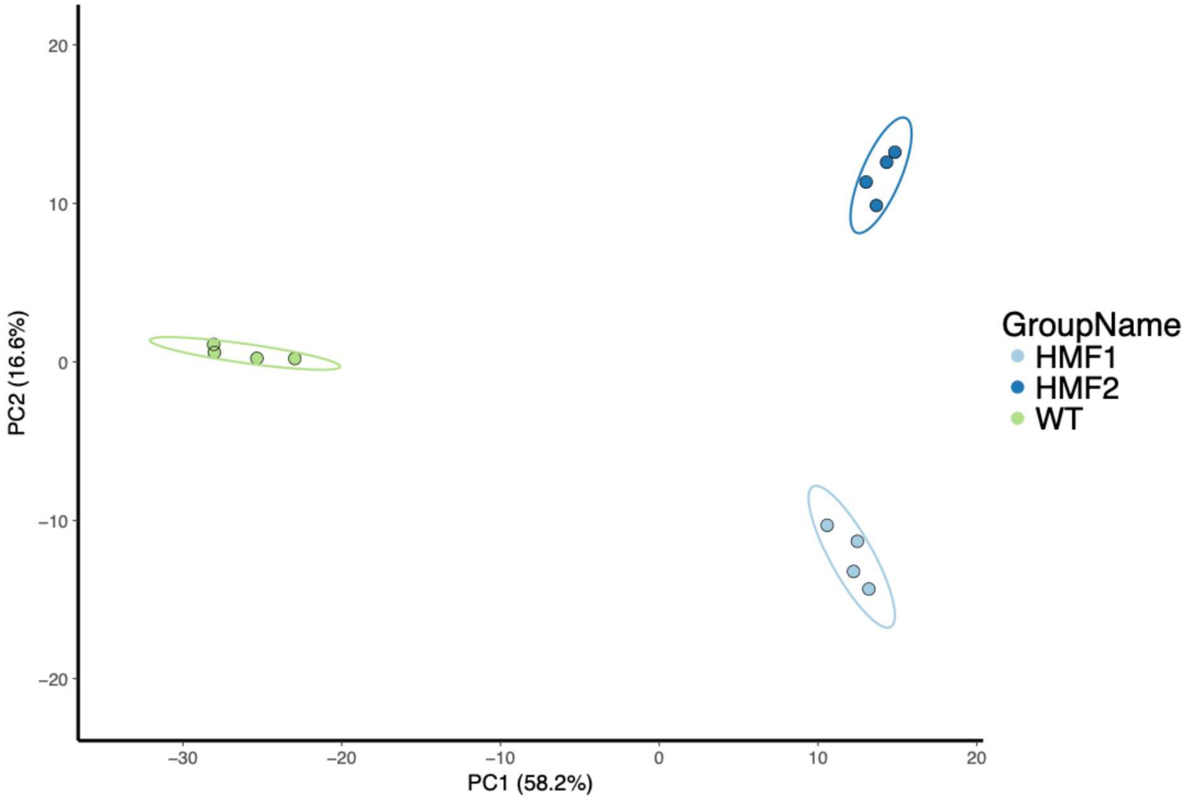


**Figure S9. Principal component analysis of shotgun lipidomics dataset in Fig 5H.**

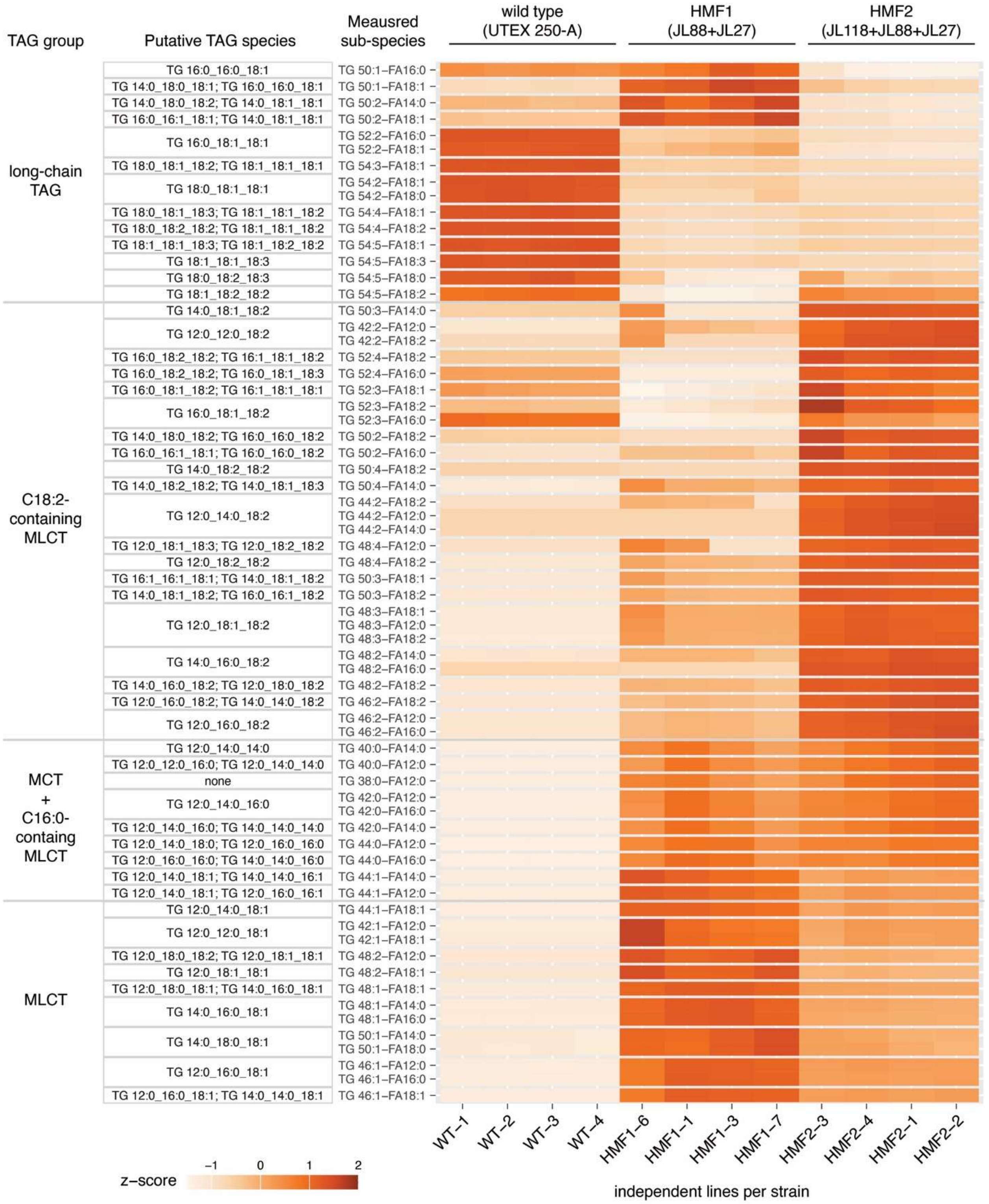

TAG group
Putative TAG species
Meausred sub-species
wild type (UTEX 250-A)
HMF1 (JL88+JL27)
HMF2 (JL118+JL88+JL27)
long-chain TAG
C18:2-containing MLCT
MCT + C16:0-containg MLCT
MLCT
TG 16:0_16:0_18:1
TG 14:0_18:0_18:1; TG 16:0_16:0_18:1
TG 14:0_18:0_18:2; TG 14:0_18:1_18:1
TG 16:0_16:1_18:1; TG 14:0_18:1_18:1
TG 16:0_18:1_18:1
TG 18:0_18:1_18:2; TG 18:1_18:1_18:1
TG 18:0_18:1_18:1
TG 18:0_18:1_18:3; TG 18:1_18:1_18:2
TG 18:0_18:2_18:2; TG 18:1_18:1_18:2
TG 18:1_18:1_18:3; TG 18:1_18:2_18:2
TG 18:1_18:1_18:3
TG 18:0_18:2_18:3
TG 18:1_18:2_18:2
TG 14:0_18:1_18:2
TG 12:0_12:0_18:2
TG 16:0_18:2_18:2; TG 16:1_18:1_18:2
TG 16:0_18:2_18:2; TG 16:0_18:1_18:3
TG 16:0_18:1_18:2; TG 16:1_18:1_18:1
TG 16:0_18:1_18:2
TG 14:0_18:0_18:2; TG 16:0_16:0_18:2
TG 16:0_16:1_18:1; TG 16:0_16:0_18:2
TG 14:0_18:2_18:2
TG 14:0_18:2_18:2; TG 14:0_18:1_18:3
TG 12:0_14:0_18:2
TG 12:0_18:1_18:3; TG 12:0_18:2_18:2
TG 12:0_18:2_18:2
TG 16:1_16:1_18:1; TG 14:0_18:1_18:2
TG 14:0_18:1_18:2; TG 16:0_16:1_18:2
TG 12:0_18:1_18:2
TG 14:0_16:0_18:2
TG 14:0_16:0_18:2; TG 12:0_18:0_18:2
TG 12:0_16:0_18:2; TG 14:0_14:0_18:2
TG 12:0_16:0_18:2
TG 12:0_14:0_14:0
TG 12:0_12:0_16:0; TG 12:0_14:0_14:0
none
TG 12:0_14:0_16:0
TG 12:0_14:0_16:0; TG 14:0_14:0_14:0
TG 12:0_14:0_18:0; TG 12:0_16:0_16:0
TG 12:0_16:0_16:0; TG 14:0_14:0_16:0
TG 12:0_14:0_18:1; TG 14:0_14:0_16:1
TG 12:0_14:0_18:1; TG 12:0_16:0_16:1
TG 12:0_14:0_18:1
TG 12:0_12:0_18:1
TG 12:0_18:0_18:2; TG 12:0_18:1_18:1
TG 12:0_18:1_18:1
TG 12:0_18:0_18:1; TG 14:0_16:0_18:1
TG 14:0_16:0_18:1
TG 14:0_18:0_18:1
TG 12:0_16:0_18:1
TG 12:0_16:0_18:1; TG 14:0_14:0_18:1
TG 50:1–FA16:0
TG 50:1–FA18:1
TG 50:2–FA14:0
TG 50:2–FA18:1
TG 52:2–FA16:0
TG 52:2–FA18:1
TG 54:3–FA18:1
TG 54:2–FA18:1
TG 54:2–FA18:0
TG 54:4–FA18:1
TG 54:4–FA18:2
TG 54:5–FA18:1
TG 54:5–FA18:3
TG 54:5–FA18:0
TG 54:5–FA18:2
TG 50:3–FA14:0
TG 42:2–FA12:0
TG 42:2–FA18:2
TG 52:4–FA18:2
TG 52:4–FA16:0
TG 52:3–FA18:1
TG 52:3–FA18:2
TG 52:3–FA16:0
TG 50:2–FA18:2
TG 50:2–FA16:0
TG 50:4–FA18:2
TG 50:4–FA14:0
TG 44:2–FA18:2
TG 44:2–FA12:0
TG 44:2–FA14:0
TG 48:4–FA12:0
TG 48:4–FA18:2
TG 50:3–FA18:1
TG 50:3–FA18:2
TG 48:3–FA18:1
TG 48:3–FA12:0
TG 48:3–FA18:2
TG 48:2–FA14:0
TG 48:2–FA16:0
TG 48:2–FA18:2
TG 46:2–FA18:2
TG 46:2–FA12:0
TG 46:2–FA16:0
TG 40:0–FA14:0
TG 40:0–FA12:0
TG 38:0–FA12:0
TG 42:0–FA12:0
TG 42:0–FA16:0
TG 42:0–FA14:0
TG 44:0–FA12:0
TG 44:0–FA16:0
TG 44:1–FA14:0
TG 44:1–FA12:0
TG 44:1–FA18:1
TG 42:1–FA12:0
TG 42:1–FA18:1
TG 48:2–FA12:0
TG 48:2–FA18:1
TG 48:1–FA18:1
TG 48:1–FA14:0
TG 48:1–FA16:0
TG 50:1–FA14:0
TG 50:1–FA18:0
TG 46:1–FA12:0
TG 46:1–FA16:0
TG 46:1–FA18:1
WT–1
WT–2
WT–3
WT–4
HMF1–6
HMF1–1
HMF1–3
HMF1–7
HMF2–3
HMF2–4
HMF2–1
HMF2–2
z–score
–1
0
1
2
independent lines per strain

**Figure S10. Relative abundance of TAG species identified by shotgun lipidomics.** Measured abundances (nmol/$10^7$ cells) per target TAG sub-species across all samples were transformed to z-score for visualization of relative abundance.

| Strain | Transgene | Anticipated activity | Marker gene | Insertion site | Line # |
|---|---|---|---|---|---|
| UTEX 250-A | NA, control | NA | NA | none | NA |
| JLM63 | NA, control | increased 18:0 | *nptII* | *fab2A-2** | 3 |
| JLM82 | *CwFATB2* | MCFA (10:0-14:0) | *nptII* | *fab2A-2* | 3 |
| JL15 | *FAB2Tp-CwFATB2* | MCFA (10:0-14:0) | *nptII* | *fab2A-2* | 3 |
| JLM24 | NA, control | NA | *nptII* | *dao1*† | 4 |
| JL20 | *CwFATB2* | MCFA (10:0-14:0) | *nptII* | *dao1* | 4 |
| JL21 | *FAB2Tp-CwFATB2* | MCFA (10:0-14:0) | *nptII* | *dao1* | 3 |
| JL13 | *BjFATB3* | 16:0 | *nptII* | *fab2A-2* | 4 |
| JL86 | *FAB2Tp-BjFATB3* | 16:0 | *nptII* | *fab2A-2* | 3 |
| JL44 | *BjFATB3* | 16:0 | *nptII* | *dao1* | 4 |
| JL60 | *FAB2Tp-BjFATB3* | 16:0 | *nptII* | *dao1* | 5 |
| JL27 | *CrLPAAT2* | *sn*-2 16:0 | *SUC2* | *dao1* | 4 |
| JL29 | *HsAGPAT1* | *sn*-2 16:0 | *SUC2* | *dao1* | 4 |
| JL52 | *FAB2Tp-CwFATB2*<br>*BjFATB3* | MCFA (10:0-14:0)<br>16:0 | *nptII* | *amt1B-1*◊<br>*dao1* | 4 |
| JL52<br>+JL27 | *FAB2Tp-CwFATB2*<br>*BjFATB3*<br>*CrLPAAT2* | MCFA (10:0-14:0)<br>16:0<br>*sn*-2 16:0 | *nptII*<br>*SUC2* | *amt1B-1*<br>*dao1* | 3 |
| JL88 | *FAB2Tp-CwFATB2*<br>*FAB2Tp-BjFATB3* | MCFA (10:0-14:0)<br>16:0 | *nptII* | *amt1B-1*<br>*dao1* | 3 |
| JL88<br>+JL27 | *FAB2Tp-CwFATB2*<br>*FAB2Tp-BjFATB3*<br>*CrLPAAT2* | MCFA (10:0-14:0)<br>16:0<br>*sn*-2 16:0 | *nptII*<br>*SUC2* | *amt1B-1*<br>*dao1* | 4 |
| JL88<br>+JL73 | *FAB2Tp-CwFATB2*<br>*FAB2Tp-BjFATB3*<br>*CrLPAAT2* | MCFA (10:0-14:0)<br>16:0<br>*sn*-2 16:0 | *nptII*<br>*SUC2* | *amt1B-1*<br>*lpaat2-1* | 4 |
| JL116 | *AtLPCAT1* | 18:2 (n-6c) | *ptxD* | *thi4*† | 3 |
| JL116<br>+JL88<br>+JL27 | *AtLPCAT1*<br>*FAB2Tp-CwFATB2*<br>*FAB2Tp-BjFATB3*<br>*CrLPAAT2* | 18:2 (n-6c)<br>MCFA (10:0-14:0)<br>16:0<br>*sn*-2 16:0 | *ptxD*<br>*nptII*<br>*SUC2* | *thi4*<br>*amt1B-1*<br>*dao1* | 4 |
| JL117 | *AtPDCT1* | 18:2 (n-6c) | *ptxD* | *thi4* | 3 |
| JL117<br>+JL88<br>+JL27 | *AtLPDCT1*<br>*FAB2Tp-CwFATB2*<br>*FAB2Tp-BjFATB3*<br>*CrLPAAT2* | 18:2 (n-6c)<br>MCFA (10:0-14:0)<br>16:0<br>*sn*-2 16:0 | *ptxD*<br>*nptII*<br>*SUC2* | *thi4*<br>*amt1B-1*<br>*dao1* | 4 |

| | | | | | |
|---|---|---|---|---|---|
| JL105 | *ApBIP1*$_{1\text{-}39}$*-mCherry-HDEL* | ER marker | *SUC2* | *dao1* | 3 |
| JL106 | *mCherry* | control | *SUC2* | *dao1* | 3 |
| JL36 | NA, control | *lpaat2-2* knock-out | *nptII* | *lpaat2-2* | 4 |
| JL33 | NA, control | *lpaat2-1* knock-out | *SUC2* | *lpaat2-1* | 4 |
| JL83 | *CrLPAAT2-Venus* | *sn*-2 16:0 | *nptII* | *lpaat2-2* | 4 |
| JL84 | *HsAGPAT1- Venus* | *sn*-2 16:0 | *nptII* | *lpaat2-2* | 3 |
| JL83+JL33 | *CrLPAAT2- Venus* | *sn*-2 16:0<br>reduced *sn*-2 18:x | *nptII*<br>*SUC2* | *lpaat2* | 6 |
| JL84+JL33 | *HsAGPAT1- Venus* | *sn*-2 16:0<br>reduced *sn*-2 18:x | *nptII*<br>*SUC2* | *lpaat2* | 5 |
| JL105<br>+JL81 | *ApBIP1*$_{1\text{-}39}$*-mCherry-HDEL*<br>*ApLPAAT2- Venus* | ER marker<br>Fluorescence-labeled ApLPAAT2 (putative) | *nptII*<br>*SUC2* | *lpaat2-2*<br>*dao1* | 3 |
| JL105<br>+JL83 | *ApBIP1*$_{1\text{-}39}$*-mCherry-HDEL*<br>*CrLPAAT2- Venus* | ER marker<br>Fluorescence-labeled CrLPAAT2 | *nptII*<br>*SUC2* | *lpaat2-2*<br>*dao1* | 3 |
| JL105<br>+JL84 | *ApBIP1*$_{1\text{-}39}$*-mCherry-HDEL*<br>*HsAGPAT1- Venus* | ER marker<br>Fluorescence-labeled HsAGPAT1 | *nptII*<br>*SUC2* | *lpaat2-2*<br>*dao1* | 3 |

* increased 18:0 due to reduced stearyl-ACP desaturase activity
† heterozygous; neutral for growth and lipid production
◇ heterozygous; neutral for growth and lipid production due to its locus on a trisomic chromosome

**Table S1. Strains generated in this study.**
